# An amphioxus neurula stage cell atlas supports a complex scenario for the emergence of vertebrate head mesoderm

**DOI:** 10.1101/2023.09.26.559513

**Authors:** Xavier Grau-Bové, Lucie Subirana, Lydvina Meister, Anaël Soubigou, Ana Neto, Anamaria Elek, Oscar Fornas, Jose Luis Gomez-Skarmeta, Juan J. Tena, Manuel Irimia, Stéphanie Bertrand, Arnau Sebé-Pedrós, Hector Escriva

## Abstract

The emergence of new structures can often be linked to the evolution of novel cell types that follows the rewiring of developmental gene regulatory subnetworks. Vertebrates are characterized by a complex body plan compared to the other chordate clades and the question remains of whether and how the emergence of vertebrate morphological innovations can be related to the appearance of new embryonic cell populations. We already proposed, by studying mesoderm development in the cephalochordate amphioxus, a scenario for the evolution of the vertebrate head mesoderm. To further test this scenario at the cell population level, we used scRNA-seq to construct a cell atlas of the amphioxus neurula, stage at which the main mesodermal compartments are specified. Our data allowed us to confirm the presence of a prechordal-plate like territory in amphioxus, and shows that cell populations of the anteriormost somites and of the ventral part of the somites present a transcriptomic profile supporting the homology with vertebrate cranial/pharyngeal and lateral plate mesoderm. Finally, our work provides evidence that the appearance of the specific mesodermal structures of the vertebrate head was associated to both segregation of pre-existing cell populations, and co-option of new genes for the control of myogenesis.

## Introduction

Chordates are an animal clade characterized by the presence of a notochord (in at least one stage of their life cycle)^1^ and that include vertebrates, tunicates (or urochordates), and cephalochordates (*i.e.* amphioxus). Even if tunicates are phylogenetically more closely related to vertebrates^2^ and share with them some morphological features absent in amphioxus^3^, they show developmental modalities and a genomic content and organization that have diverged considerably from the chordate ancestral state^4^. On the other hand, amphioxus exhibit relatively conserved morphological, developmental, and genomic characteristics, and represent a model of choice for studying chordate evolution and the emergence of vertebrate novelties^5,6^.

The gastrula of cephalochordates has two germ layers: the ectoderm, which forms the epidermis and the central nervous system, and the internal mesendoderm, which develops into mesodermal structures in the dorsal part, and into endodermal structures in the ventral region^7^. Unlike vertebrates, the mesoderm is first simply divided during neurulation into the axial territory forming the notochord, and the paraxial domain that becomes completely segmented into somites from the most anterior to the posterior part of the embryo. In vertebrates, in addition to the notochord and somites, the mesoderm is subdivided into other territories: the lateral plate mesoderm in the trunk that forms several structures among which part of the heart and circulatory system, blood cells, fin buds or excretory organs^8^; and the prechordal plate (axial) and cranial/pharyngeal (paraxial, unsegmented) mesoderm in the anterior region that form head muscles and part of the heart^9,10^. If we consider that the amphioxus mesoderm organization could resemble that of the chordate ancestor, these mesodermal territories represent vertebrate specific traits that contributed to the acquisition of particular structures, including the complex vertebrate head.

Based on previous work, we have proposed a multi-step scenario for the evolution of the vertebrate anterior mesoderm^11,12^. The first step consists in the segregation of the ventral mesoderm from the paraxial mesoderm and loss of its segmentation. This implies that the ventral part of amphioxus somites is homologous to the vertebrate lateral plate mesoderm. The second step corresponds to the loss of the paraxial mesoderm in the anterior part of the embryo. This would have enabled the relaxation of the developmental constraints imposed by the anterior somites, and the remodelling of the axial and lateral plate mesoderm resulting in the appearance of the prechordal plate and cranial/pharyngeal mesoderm. This would mean that i) the cranial/pharyngeal mesoderm has a lateral rather than paraxial origin, and partly shares a common developmental program with the amphioxus anterior somites and ventral part of posterior somites, and ii) the prechordal plate is in part homologous to the amphioxus anterior notochord.

Here we sought to explore the evolutionary origin of the vertebrate head mesoderm from a cell type perspective. In order to compare embryonic cell types between amphioxus and vertebrates, we conducted a scRNA-seq analysis of the *Branchiostoma lanceolatum* neurula (N3)^13,14^. The neurula stage shows the highest global transcriptional similarity with vertebrates^15^, corresponding to the chordate phylotypic stage ^16^, and our cell atlas uncovers the gene expression signatures of most of the previously described embryonic territories at this stage. Concerning the mesoderm compartment, we found a cell population with a mixed profile between endoderm and notochord, supporting the existence of a transient prechordal plate-like structure in amphioxus^12,17^. We also show that the first somite pair cells form a population with a transcriptomic profile different from the posterior somites, highlighting the peculiarity of this somitic pair. Moreover, these cells express orthologues of vertebrate genes expressed in both head and lateral plate mesoderm and their derivatives, bringing further support to our evolutionary scenario, and suggesting how, from pre-existing cell populations, new embryonic territories might have emerged in vertebrate anterior mesoderm. Finally, the functional study in transgenic zebrafish lines of regulatory regions of *Gata1/2/3*, *Tbx1/10* and *Pitx* also supports the lateral origin of the cranial/pharyngeal mesoderm and gives insights into how genes that were presumably not controlling muscle formation in the chordate ancestor were co-opted as master genes of the myogenesis program in the vertebrate head.

## Results and discussion

### A cell atlas of the amphioxus neurula stage embryo

To build a transcriptional cell atlas of the amphioxus neurula stage (N3), we applied MARS-seq^18^ to embryos at 21 hpf (hours post-fertilization, at 19°C) (**Fig. 1a**). Briefly, cells were dissociated and alive single cells (calcein positive, propidium-iodide negative) were sorted into 384-well plates, followed by scRNA-seq library preparation. At this developmental stage, the embryo is made of around 3,000 cells and we sampled in total 14,586 single-cell transcriptomes, representing approximately a five-fold coverage. These cells were grouped into 176 transcriptionally coherent clusters (referred to as “metacells”^19^ (**Fig. 1b, Supplementary Fig. 1a**). Metacells were further assigned to a tissue/cell type by using transcriptional signatures of known marker genes: epidermis, endoderm, mesoderm, muscular somite and neural (**Fig. 1c**). The proportion of cells assigned to each structure/germ layer was overall consistent with cell counting in 3D embryos reconstructed using confocal imaging of labelled nuclei followed by image segmentation, with more than half of the cells belonging to the epidermis (**Fig. 1d**).

**Figure 1.**
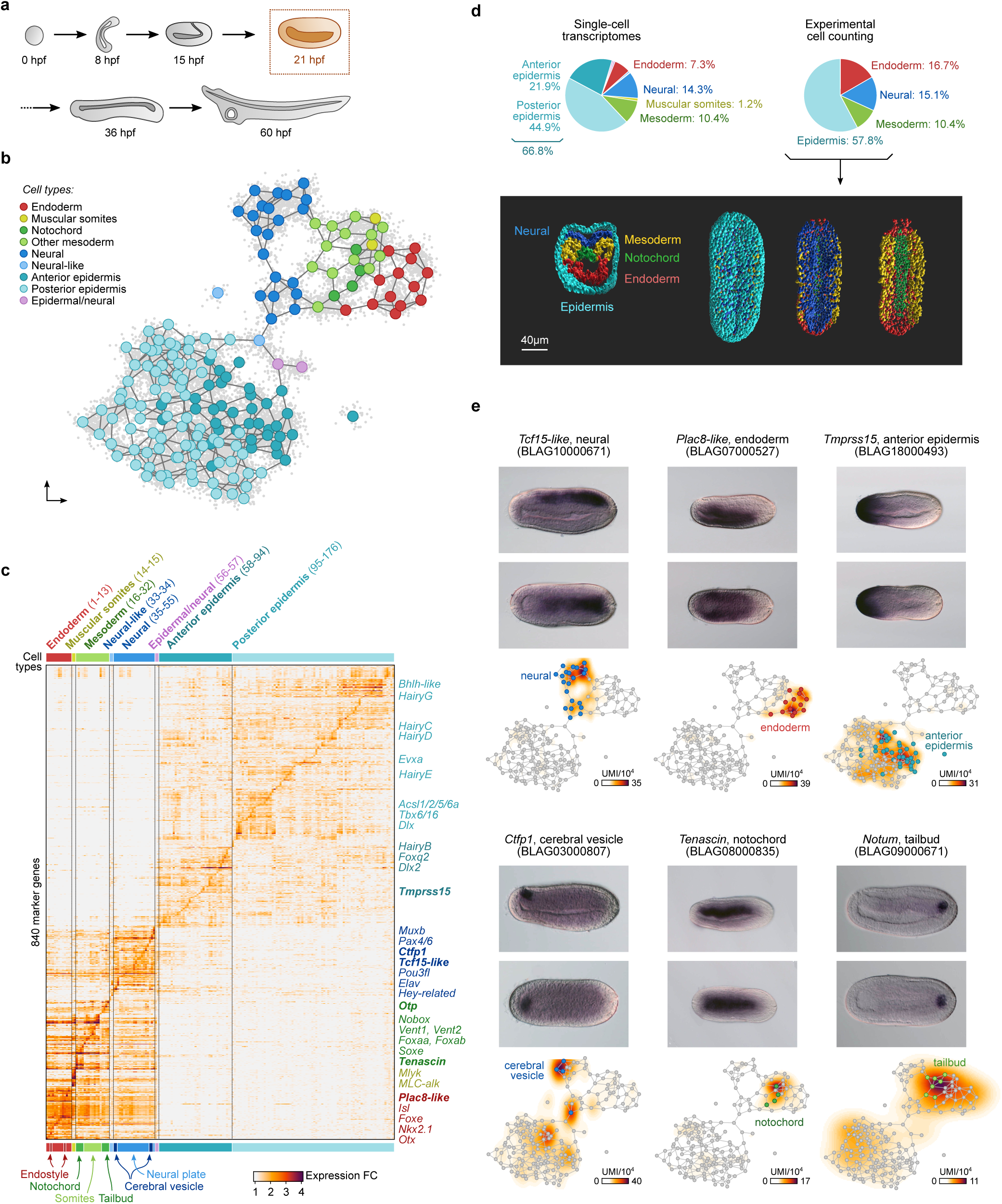
Amphioxus neurula cell type atlas. **a,** drawings of Mediterranean amphioxus developmental stages from the egg to the larva (with one open gill slit) stage. The developmental time (hours post fertilization, hpf) is given for embryos raised at 19°C, and we highlight the neurula stage presented in this study (21 hpf). **b**, Two-dimensional projection of cell clusters (metacells) using a force-directed layout based on the co-clustering graphs for individual cells (see *Methods*). Metacells are colour-coded by cell type. **c,** Normalized fold change expression of top variable genes (rows) per metacell (columns, grouped by cell type). For each metacell, we selected up to 30 markers with a minimum fold change ≥ 2. Selected gene names from known markers, used to annotate each cell type, are indicated to the right of the heatmap. Genes in bold case are shown in panel d. **d,**. Pie charts depicting the fraction of cells mapped to each cell type among the cell transcriptomes and the cell counting experiment (top); and the 3D reconstruction with assignment of nuclei to each germ layer (bottom). A transverse section is shown on the left, and dorsal views with anterior to the top on the right (full, without epidermis nuclei, without epidermis and neural cells nuclei. **e,** Expression profile of previously unknown marker genes for specific cell types (neural, endoderm, anterior epidermis, cerebral vesicle, notochord, and tailbud) analyzed by *in situ* hybridization (ISH, top, with anterior to the left and dorsal to the top in side views) and corresponding two-dimensional expression maps (bottom, based on the same layout as panel b). Gene expression is shown as density maps representing UMI counts (per 10,000 UMIs) in each cell.

Gene expression signatures across epidermal metacells shows that this tissue is not homogenous. For example, we recognized anterior epidermal cells (i.e. expressing *Arpd2, Fgfrl*, *Fzd5/8*, *Pax4/6*)^20–23^, posterior cells (*Cdx*, *Tbx6/16*, *Wnt3*)^24–26^ and subpopulations of potential epidermal sensory cells (*Delta*, *Elav*, *Tlx*)^27–29^. Among the neural metacells, we identified several metacells corresponding to the cerebral vesicle (anterior central nervous system, *Otx*^30^). Concerning the mesodermal cell populations, we could assign several metacells to the notochord (*Cola*, *Foxaa*, *Mnx*, *Netrin*)^31–34^ and tailbud compartments (*Nanos*, *Piwil1*, *Vasa*, *Wnt1*)^26,35,36^. In the endoderm, one metacell could be assigned to the ventral endodermal region that later develops into the endostyle and the club-shaped gland (*Foxe*, *Nkx2.5*)^37,38^. We further validated our atlas by analysing by *in situ* hybridisation the expression of several genes with undescribed patterns, including genes enriched in neural plate (*Tcf15-like*), endoderm (*PLAC8 motif-containing protein 1)*, anterior epidermal (*ST14-like*), cerebral vesicle (*Calcitonin Family Peptide 1* (*Ctfp1*)), notochord (*Tenascin*) or tailbud (*Notum*) populations (**Fig. 1e and Supplementary Fig. 2**). Overall, our single-cell transcriptomic atlas uncovers the diversity of cell states associated to each major germ layer in the amphioxus neurula.

### Cross-species comparison of neurula stage embryonic tissues

To gain insights into the evolutionary affinities of amphioxus neurula stage tissues, we compared aggregated expression profiles of the different structures and tissues with those of three other chordates, using published developmental single-cell atlases for *Ciona intestinalis*^39^, *Xenopus laevis*^40^ and *Danio rerio*^41^ (**Fig. 2a**). We focused our comparative analysis on stages approximately corresponding to the amphioxus neurula stage^15^ and used single-cell expression profiles similarly grouped into embryonic tissues.

**Figure 2.**
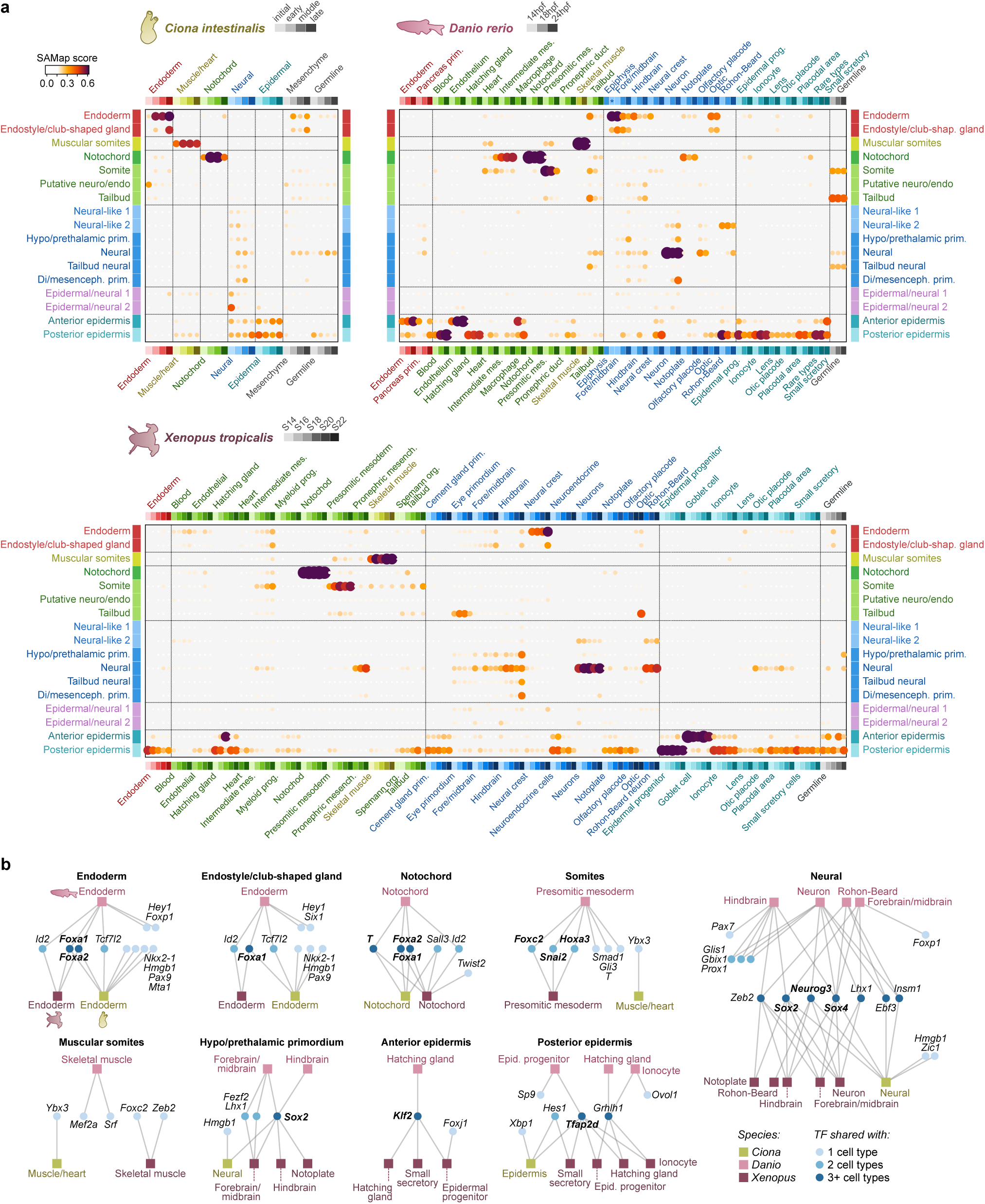
Cross-species comparison with other chordate developmental datasets. **a,** Comparison between cell type transcriptomes of the amphioxus neurula stage (rows) and matched developmental time-points (columns) in the chordates *Ciona intestinalis* (initial to late tailbud stage), *Danio rerio* (14 hpf to 24 hpf), and *Xenopus tropicalis* (S14 to S22 stages). Cell type similarity was measured using SAMap scores based on all available pairwise markers (see *Methods*). Cell types are colour-coded by developmental layer (endoderm, mesoderm/muscle, neuroectoderm, ectoderm, and other), and, in the case of the multi-stage chordate datasets, by developmental time-point (colour intensity). **b,** Graph representation of transcription factors (TFs, circular nodes) shared (i.e. connected by an edge) between amphioxus and homologous cell types in *Ciona*, *Danio* and *Xenopus* (square nodes). Specific TFs are considered to be shared between two cell types if they are significantly overexpressed in both. TFs in bold are shared between all species considered. For amphioxus, we required fold-change > 1.25, and BH-adjusted *p-*value < 0.05. For the matched cell types from other species, we required significant overexpression in at least one of the developmental time-points considered. A complete list of genes shared between all pairs of cell types is available in Supplementary Table 1.

In all three pairwise comparisons, the notochord showed the strongest transcriptional similarity and shared expression of the TFs *Brachyury2* (*T)*, *Foxaa* and *Foxab* (*Foxa1* and *Foxa2)* (**Fig. 2b**). Likewise, amphioxus differentiated muscular somites resemble muscle/skeletal muscle in tunicates and both vertebrates (**Fig. 2a**), albeit with different sets of TFs between species (**Fig. 2b**). In contrast, the non-muscular part of amphioxus somites resembles vertebrate presomitic mesoderm and shares expression of the TFs *Foxc* (*Foxc2)*, *Snail* (*Snai2)* and *Hox3* (*Hoxa3)* (**Fig. 2b**). Amphioxus neural cells also resemble vertebrate neural populations and co-express neural TFs like *Soxb1c* (*Sox2), Soxc (Sox4)* and *Neurogenin* (*Neurog3)*. These same TFs are also shared by tunicate neural cells, but the overall transcriptome does not show similarity with amphioxus neurons. The opposite is true for the endodermal transcriptome: amphioxus endoderm transcriptome matches that of tunicates, but not vertebrate endoderm, although the TFs *Foxaa* and *Foxab* (*Foxa1* and *Foxa2)* are expressed in all of them (**Fig. 2b**). Finally, the amphioxus anterior epidermis looks more distinct than the posterior one. Among vertebrate epidermal cells, its most similar pairs are secretory cells both in *Danio* and *Xenopus* (termed “Goblet cells” there). But it also hits different mesodermal tissues in *Danio* (e.g. the endothelium). On the other hand, the amphioxus posterior epidermis is broadly similar to many epidermal cell types of the two vertebrates, most notably the epidermal progenitors and ionocytes.

When examining the lists of shared markers between transcriptionally similar embryonic tissues/cell types (**Supplementary Table 1**), we observed a general overrepresentation of transcription factors (TFs) and chromatin factors compared with effector genes, as expected when comparing undifferentiated cell populations.

### The accessible chromatin landscape of amphioxus neurula stage

To interrogate the regulatory logic underlying the observed cell-specific transcriptomes, we performed bulk ATAC-seq experiments in neurula stage embryos. We defined a total of 51,028 ATAC-seq peaks and assigned them by proximity to 19,069 genes (median 2,05 peaks per expressed gene) (**Supplementary Fig. 1b-i)**. We then grouped these peaks according to the expression pattern of the associated genes and conducted motif enrichment analysis on these regulatory element groups, using a combination of *de novo* inferred and known motifs (see Methods). This analysis revealed 317 distinct motifs with significant enrichments in specific cell populations (**Fig. 3a**).

**Figure 3.**
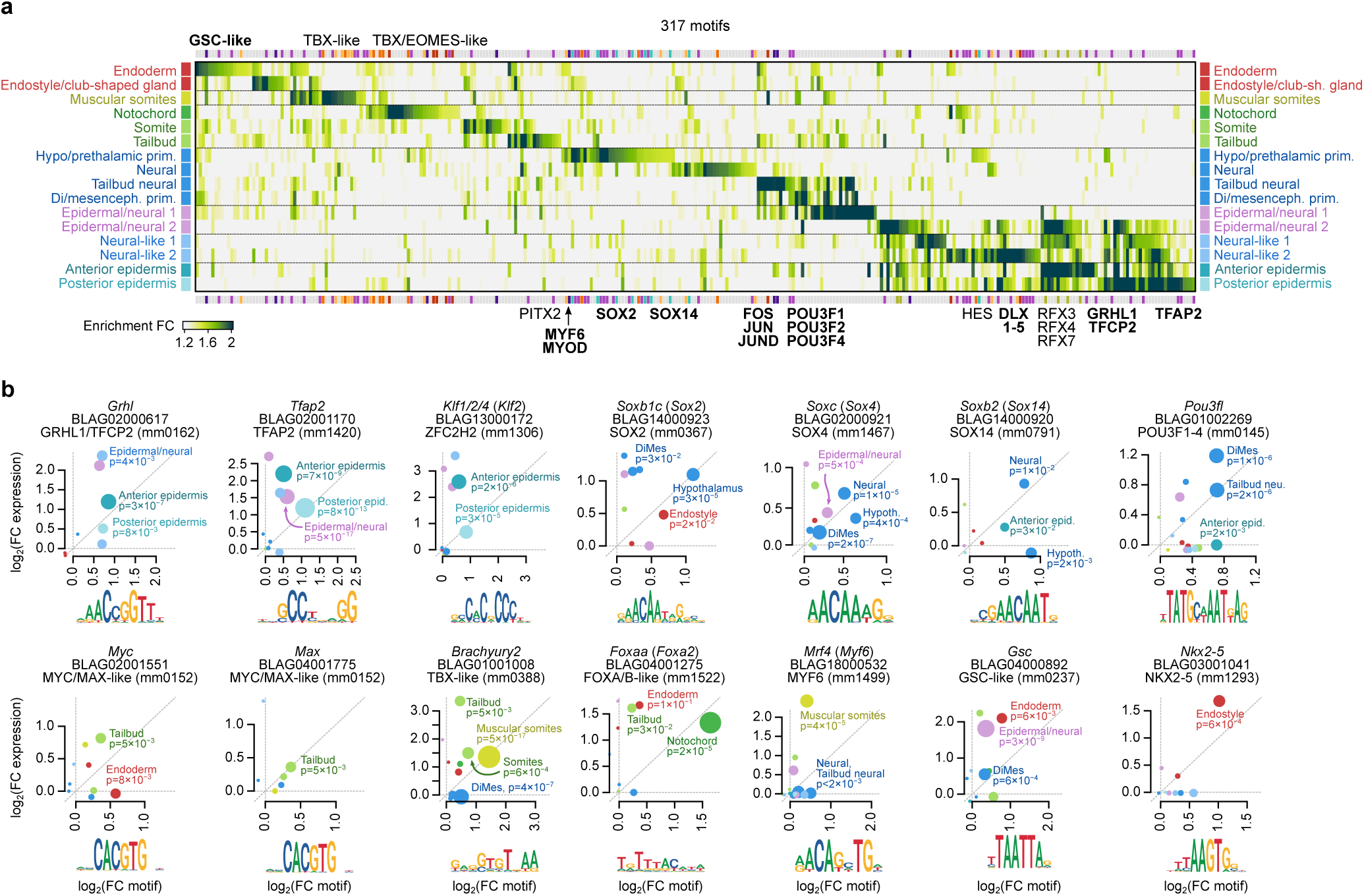
Regulatory landscape of neurula cell types. **a,** Heatmap representing the enrichment of specific TF binding motifs (columns) in the regulatory regions of genes in each cell type of the amphioxus neurula (rows). Names from selected motifs are indicated next to the heatmap (in bold those that also appear in panel b). The amphioxus motif library was obtained by merging experimentally determined vertebrate motifs from CIS-BP with *de novo* inferred motifs for each amphioxus cell type, and removing redundancy (see *Methods*). Therefore, motif names do not represent specific amphioxus TFs, but rather sequence similarity with motifs of vertebrate homologs. The regulatory regions associated with each gene were obtained from a bulk ATAC-seq experiment. **b,** Examples of cell type-specific amphioxus TFs whose expression levels (vertical axis, as log_2_(FC)) match the enrichment of associated motifs (horizontal axis; shown below as information content logos). Circle size is proportional to the BH-adjusted −log_10_(*p*) of motif enrichment (shown only for significant enrichment at *p* < 0.01).

The identified motifs are consistent with known TF regulators in amphioxus and other metazoans and, in addition, motif enrichments often parallel the expression of the associated TFs (**Fig. 3b, Supplementary Fig. 3**). For example, in peaks assigned to epidermal genes, we found enrichment for motifs like Dlx, Grhl, Klf1/2/4, Rfx1/2/3 or Tfap2, coincident with the expression of *Dlx*, *Klf1/2/4* and *Tfap2* in epidermal metacells (**Fig. 3b**). Interestingly, these TFs are part of the *in silico* reconstructed gene regulatory network controlling epidermis development described in amphioxus^42^ and are known epidermal fate determinants in vertebrates^43–46^ (**Fig. 2b**).

In endodermal cells, we found a slight but non-significant enrichment of a Fox motif in the regulatory regions of endodermal marker genes, possibly linked to the expression of *Foxaa* and *Foxab* in these tissues (**Supplementary Fig. 3**). Furthermore, in the endoderm and the endostyle, we also observed the coincident expression/motif enrichment of *Gsc* and *Nkx2-5/6*, respectively (**Fig. 3b**).

Neural cell types exhibited expression of various Sox and Pou family TFs and concomitant enrichment of their associated motifs, including *Soxc* (*Sox4*) and *Soxb2* (*Sox14*) in multiple neural tissues, the hypothalamus-specific expression/enrichment of *Soxb1c* (*Sox2)*, and *Pou3fl* (*Pou3f4)* in the Di-Mesencephalic primordium^20^ and neural tailbud cells (**Fig. 3b**). The neural specificities of these TFs appear to be conserved across vertebrates (**Fig. 2b**) and SoxB1 and Pou3f family factors have been proposed as potential major regulators of nervous system development in amphioxus^42^. The activity of *Pou3fl* (*Pou3f4)* in neural tailbud cells is also consistent with the function of TFs from these families in stemness maintaining in vertebrates^47^. This is also the case for the *Myc*/*Max* HLHs in the non-neural tailbud population, as observed in mouse^48^.

The peaks associated to genes overexpressed in non-muscular somite cell populations are enriched in T-box motifs, consistently with the expression in our dataset of various TFs of this family such as *Eomes/Tbr1/Tbx21*, *Tbx15/18/22,* and *Brachyury2* (**Fig. 3b**) and with previously reported expression of these genes in forming somites^49–51^. The muscular somite population peaks are enriched in motifs shared with the non-muscular somite, but are also enriched in motifs for Myogenic Regulatory Factors (MRF) such as *Mrf4* (*Myf6)*, which is also highly expressed in this cell type (**Fig. 3b**), in line with both the expression of the various amphioxus MRFs described by *in situ* hybridization^52^, and the role of their orthologues in vertebrate myogenesis^53^. The strongest TF-motif association concerns the previously reported notochordal marker *Foxaa* (*Foxa2)* (**Fig. 3b**)^34^, which is also shared with tunicates and vertebrates in our cross-species cell type comparisons (**Fig. 2b**). Overall, the accessible chromatin landscape of the neurula stage revealed the regulatory motif lexicons underlying amphioxus embryonic cell identities.

### Characterization of neural, endodermal and somitic cell populations

We then focused on the detailed analysis of specific cell populations. To this end, we performed separate clustering of single cells classified as belonging to the neural tissue (*sensu stricto*, derived from the neural plate), to the endoderm and to the somites (muscular and non-muscular).

Neural plate cells could be clustered into 22 metacells (**Fig. 4a, b, Supplementary Fig. 2 and 4**). Among these, we recognized three metacells corresponding to the cerebral vesicle (4, 12, and 22). Metacells 4 and 22 coexpress the known marker genes *Arpd2*, *Fezf, Fgfrl, Fgf8/17/18,* and *Otx*^20,21,30,54^ (**Supplementary Fig. 4**) together with the newly described genes *Celf3/4/5/6* (**Fig. 4a, b**) and *Ctfp1* (**Fig. 1e, Supplementary Fig. 4**) and correspond to the Hypothalamo-prethalamic primordium as previously described^20^ with metacell 4 overexpressing *Six3/6*^20^ (**Supplementary Fig. 4**) and hence representing its rostral part. On the other hand, metacell 12 shows expression of *Otx* and *Pax4/6,* a combination typical of the Di-Mesencephalic primordium^20^ (**Supplementary Fig. 4**). Metacells 20 and 21 co-express the posterior gene markers *Cdx*, *Nanos*, *Vasa* and *Wnt1*^25,35,55^ (**Supplementary Fig. 4**), together with *Bolla, Otp* and *Zf-Ring Protein* described here (**Fig. 4a, b, Supplementary Figure 2 and 4**), suggesting that these metacells represent the posterior-most neural plate. In addition to expressing posterior markers, metacell 9 also expresses *Netrin* that marks the floorplate^33^ (**Supplementary Fig. 4**). According to the expression of the floor plate marker genes *Chordin*, *Foxaa*, *Goosecoid*, *Netrin*, *Nkx2.1* and *Nkx6*^20,26,33,34,56–59^ (**Supplementary Fig. 4**), metacells 8 and 14 could be assigned to this structure, with metacell 8 additionally expressing the posterior genes *Cdx* and *Hox3*^20,25,60^ (**Supplementary Fig. 4**). The expression of *Msx*, *Pax3/7* and *Snail*^20,61–63^ in metacells 11 and 13 indicate they belong to the neural plate border with metacell 13 expressing the anterior marker *Gremlin*^64^, and metacell 11 expressing the posterior gene *Hox3*^20,60^ (**Supplementary Fig. 4a**). We could also recognize metacells 2, 7 and 10 as segmentally arranged neurons co-expressing *Islet*^65^ (**Supplementary Fig. 4**) and *Nhlh1/2* (**Fig. 4a,b**), with metacell 10 corresponding to a specific pair of neurons characterized by *Celf3/4/5/6* and *Igfbp* expression (**Fig. 4a,b**). All the other metacells show few specific markers and could represent differentiating cells. These cells express different combinations of the known neural genes *Elav*^27^ and *Neurogenin*^66^, together with *Hey-related* (**Fig. 4b**), *Prox* (**Supplementary Fig. 2, 4**) and *Tcf15-like* genes (**Fig. 1e, Supplementary Fig. 4**).

**Figure 4.**
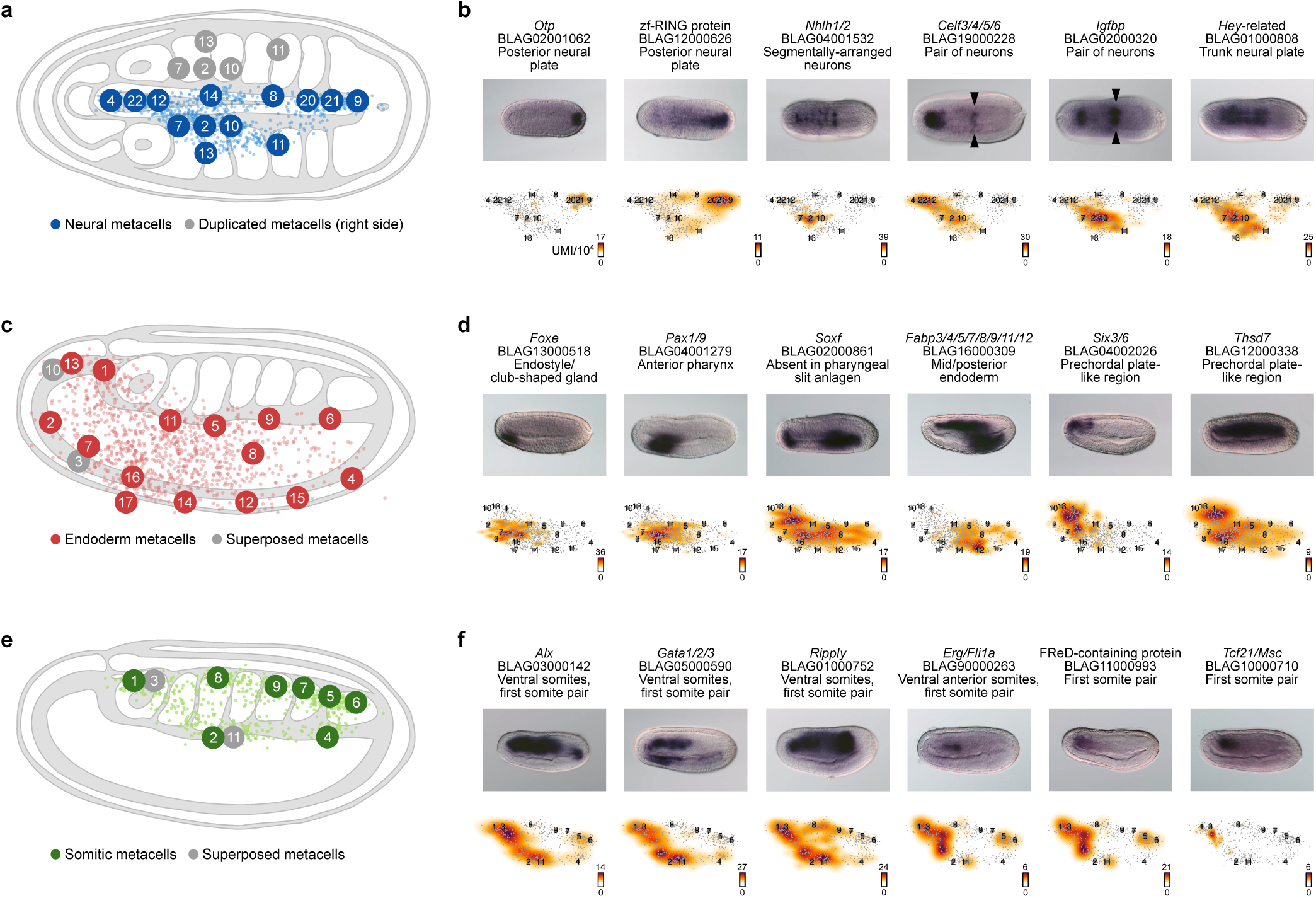
Subclustering reveals new cell types. **a,** 2D projection of neural metacells on a dorsal view scheme of an amphioxus neurula stage embryo with anterior to the left. **b**, Gene expression distribution on 2D projected cells for selected neural gene markers and corresponding ISH. Gene expression is shown as density maps representing UMI counts (per 10,000 UMIs) in each cell, with cells positioned in the vicinity of their corresponding metacells. **c**, 2D projection of endoderm metacells on a side view scheme of an amphioxus neurula stage embryo with anterior to the left and dorsal to the top. **d**, Gene expression distribution on 2D projected cells for selected endoderm gene markers and corresponding ISH. **e**, 2D projection of somite metacells on a side view scheme of an amphioxus neurula stage embryo with anterior to the left and dorsal to the top. **f**, Gene expression distribution on 2D projected cells for selected somite gene markers and corresponding ISH.

Concerning the endodermal compartment, we could recognize metacells corresponding to the main known territories (**Fig. 4c, d and Supplementary Fig. 2 and 5**). The expression of the ventral marker *Nkx2.1*^67^, together with anteriorly expressed genes such as *Dmbx*, *Fgfrl*, *Fzd5/8* and *Sfrp1/2/5*^20,21,23,59,68,69^ (**Supplementary Fig. 5**) indicates that metacell 2 corresponds to the ventral anterior endoderm territory whereas metacells 3 and 7 show a combination of marker genes that are typical of the ventral endoderm that later develops into the club-shaped gland and the endostyle such as *Foxe*, *Nkx2.5*, *Tbx1/10* and *Pax1/9*^37,38,70,71^ (**Fig. 4c, d and Supplementary Fig. 5**). The expression of *Pitx* in metacell 3 suggests that metacells 3 and 7 correspond to the left and right part of this territory, respectively^72^. Posterior to that, metacells 16 and 17 that are characterized by low or no expression of *Soxf* correspond to the first pharyngeal slit anlagen^73^ while metacell 14 expresses both *Irxc* and *Foxaa*, a combination specifically observed in a region that is just behind it^34,74^ (**Fig. 4c, d and Supplementary Fig. 5**). Metacells 5 and 11 express *Pax1/9* but no ventral markers and could correspond to the dorsal mid endoderm region^70^ (**Fig. 4c,d**). Metacells 8 and 12 show a very similar profile with an enrichment in transcripts of mid/posterior endoderm markers such as *Nkx2.2*, *Foxaa*^34,75^ (**Supplementary Fig. 5**), and the newly described gene *Fabp3/4/5/7/8/9/11/12* (**Fig. 4c, d**) with metacell 12 additionally expressing *Gata4/5/6* indicating that the corresponding cells are more ventral than those from metacell 8^76^ (**Supplementary Fig. 5**). Metacell 9 has a transcriptional profile similar to that of metacells 8 and 12 combining expression of the mid/posterior marker *Foxaa*^34^ (**Supplementary Fig. 5**) and absence of *Pax1/9* expression^70^ (**Fig. 4c, d**). The posterior marker *Wnt8*^55^ is expressed in metacells 4 and 6 with metacell 4 also expressing the ventral marker *Gata4/5/6*^76^, and, hence, representing the ventral posterior territory (**Supplementary Fig. 5**). Finally, metacells 1, 10 and 13 are characterized by an enrichment in anterior markers *Dmbx*, *Fgfrl*, *Fzd5/8* and *Sfrp1/2/5*^21,23,59,68,69^ as well as *Six3/6*, *Six4/5* and *Zic*^77,78^ (**Fig. 4c, d and Supplementary Fig. 5**). They show a transcriptional signature of the anterior dorsal mesendoderm, a region which is continuous with the notochord *per se* posteriorly, and which is continuous laterally with the endoderm *per se*. Metacell 1 is also expressing the newly described gene *Thsd7* (**Fig. 4c,d**), together with *Brachyury2*, *Pax3/7* and *Zeb*^42,51,61^ and lacks *Nkx2.1* expression^58^ (**Supplementary Fig. 5**) suggesting it represents the axial part of this region, whereas metacells 10 and 13, expressing *Nkx2.1*, would correspond to the paraxial more ventral portion that latter form the left and right Hatschek’s diverticula^58^ (**Supplementary Fig. 5**). Therefore, metacell 1 represents a potential prechordal plate-like territory showing a transcriptomic profile characterized by anterior and axial markers together with endodermal markers. Such a territory was already proposed to exist in amphioxus based on both cell behaviour and gene expression of several marker genes^12,17,79^ but our data highlight the strong difference in its transcriptomic profile compared to the other notochord cells, reinforcing the idea that ancestral chordates possessed a prechordal plate-like region that later evolved specific functions in vertebrates.

Finally, re-clustering of cells assigned to the somites resulted in 12 metacells (**Fig. 4e, f and Supplementary Fig. 2 and 6**). As expected, we found a population (metacell 8) with a profile typical of the muscular part of somites that starts to differentiate, characterized by the expression of *Mef2*, *Lmo4*, several MRFs, together with *MLC-alk*^52,80,81^ (**Supplementary Fig. 6**) and the newly described gene *Titin-like* (**Supplementary Fig. 2 and 6**). Metacells 7 and 9 have similar profiles and also express *Titin-like* and several MRFs^52^ (**Supplementary Fig. 2 and 6)** together with *Brachyury2*, *Delta*^29,51^ and the newly described gene *Twist-like* (**Supplementary Fig. 2 and 6**). They hence correspond to the last somites that have just been formed, with metacell 7 more posterior as indicated by the expression of *Wnt1* or *Wnt4*^55^. More posteriorly, metacell 5 is characterized by the expression of newly described tailbud gene markers such as *Bicc*, *Otp*, *SF2 family helicase* (**Fig. 4b, Supplementary Fig. 2 and 6**), together with *Vasa*, *Nanos* and *Wnt1*, *4* and *6*^36,55^ but also expresses *Brachyury2* and *Mrf4*, a combination corresponding to the tailbud somitic part^51,52^ (**Supplementary Fig. 6**). Metacells 4 and 6 also express tailbud markers but do not express MRF genes. Moreover, metacell 4 is characterized by an enrichment in transcripts of the ventral markers *Gata1/2/3* and *Vent1/Vent2*^76,82,83^ (**Fig. 4e, f and Supplementary Fig. 6**). The most important novelty concerns the first somite pair, which clearly shows a transcriptomic profile divergent from the other pairs. Metacells 1 and 3 correspond to this first pair, with metacell 1 representing the right somite, and metacell 3 the left one (**Fig. 4e, f**). Indeed, contrary to metacell 1, cells of the latter express the left side marker *Pitx*^72^ as well as *Gremlin*, which is expressed in the first left somite at this stage^64^ (**Supplementary Fig. 6**). Both metacells express the anterior marker *Fgfrl*^21^ (**Supplementary Fig. 6**), and three newly described markers: *Erg/Fli1a*, *Tcf21/Msc* and *FReD containing protein* (**Fig. 4e, f**). They also express the ventral somite marker genes *Alx*, *Gata1/2/3*, *Ripply* and *Vent1/Vent2*^32,76,82–84^ (**Fig. 4e, f and Supplementary Fig. 6**). To note, no Wnt genes are expressed in these metacells, whereas the ventral markers are expressed together with *Wnt16*^55^ in metacells 2 and 11 that correspond to the ventral region of the formed somites posterior the the first pair (**Supplementary Fig. 6**). Interestingly, *Erg/Fli1a* is orthologous to *Fli-1* which is implicated in vertebrate hemangioblast development together with *Vegfr* and *Scl/Tal-1*^85^. The amphioxus orthologues of the latest were also shown to be expressed in the first somite pair^76^, reinforcing the proposition of homology between this first pair of somites and the embryonic hematopoietic/angiogenic field of vertebrates that derives from the lateral plate mesoderm. On the other hand, *Tcf21/Msc* is orthologous to *Tcf21/Capsulin* and *Msc/MyoR* that are main regulators of head muscle myogenesis in vertebrates, upstream of MRFs^86–88^, suggesting that the first somite pair of amphioxus has a profile that resembles both vertebrate head and lateral plate mesoderm.

### The evolution of the chordate anterior mesoderm

The most striking feature of the amphioxus neurula highlighted by our data is the presence of three cell populations with a peculiar transcriptional profile: cells of the first left and right somites (metacells 1 and 3, **Fig. 4e, f**), and cells that could correspond to a prechordal plate-like structure (metacell 1, **Fig. 4c, d**). The first somite pair in amphioxus has long been proposed as being distinct from the other pairs, and we previously showed that this somite pair is the only one whose formation is controlled by the FGF signalling pathway^11,12,54^. Our molecular atlas additionally shows that the cells of the first pair of somites transcriptionally resemble vertebrate head and lateral plate mesoderm (metacells 1 and 3, **Fig. 4e, f and Supplementary Fig. 5**), while the cells of the ventral part of amphioxus somites posterior to the first pair express orthologues of genes expressed in vertebrates lateral plate mesoderm or derivatives (metacells 2 and 11, **Fig. 4e, f and Supplementary Fig. 5**). These data support the homology we proposed between vertebrate lateral plate mesoderm and amphioxus ventral part of the somites as well as the ventral origin of vertebrate cranial/pharyngeal mesoderm. Such proposed homology based on transcriptomic profile should reflect a conserved regulatory logic. Considering homology at the gene expression regulation level, we reasoned that if our scenario for vertebrate mesoderm evolution supported by our cell atlas is correct, regulatory regions of genes that are active at the neurula stage in amphioxus in the ventral region of somites could drive expression of a reporter gene in the vertebrate lateral plate and head mesoderm, as a reminiscence of an ancestrally shared regulatory program. Among such genes, *Gata1/2/3* is the transcription factor with the highest enrichment (fold change) in metacells 2 and 11 of the somite reclustering analysis (**Fig. 4e, f**), which we could assign to the ventral region of the somites. We decided to test whether the regulatory elements controlling the expression of amphioxus *Gata1/2/3* at this stage are recognized by any tissue/cell type specific regulatory state in zebrafish, which would point at evolutionary conservation (at least partially) of *Gata1/2/3* regulation. To this end, we generated transgenic reporter constructs for four putative regulatory regions selected using ATAC-seq data (**Fig. 5a**). We tested the heterologous activity of these regions by generating F0 transgenic zebrafish. The transcription factor binding motif composition of the four tested regions differs completely (**Supplementary Table 2**), which means that we should expect distinct activities when separately exposed to zebrafish regulatory states. Only one region was able to drive the reporter gene expression (*eGFP*) in a restricted manner in F0 embryos and we generated F1 transgenics for the corresponding construct. We observed green fluorescence in the head mesoderm at 24 hpf and in both the pectoral fin buds and the head mesoderm at 48 hpf (**Fig. 5b**). The genomic sequence cloned in this reporter assay contains motifs for *Alx* and *Foxc1* (**Fig. 5a, Supplementary Table 2**) and both *Alx1* and *Foxc1a* are expressed in the head mesoderm of zebrafish^89–91^. It also contains motifs for *Prrx1* (**Fig. 5a, Supplementary Table 2**), with *Prrx1a* and *Prrx1b* being expressed in the zebrafish head mesoderm, branchial arches and pectoral fin buds^92^, suggesting that part of the factors that control gene expression of *Gata/1/2/3* in the ventral part of amphioxus somites also regulate the expression of genes in both the head and lateral plate mesoderm in vertebrates.

**Figure 5.**
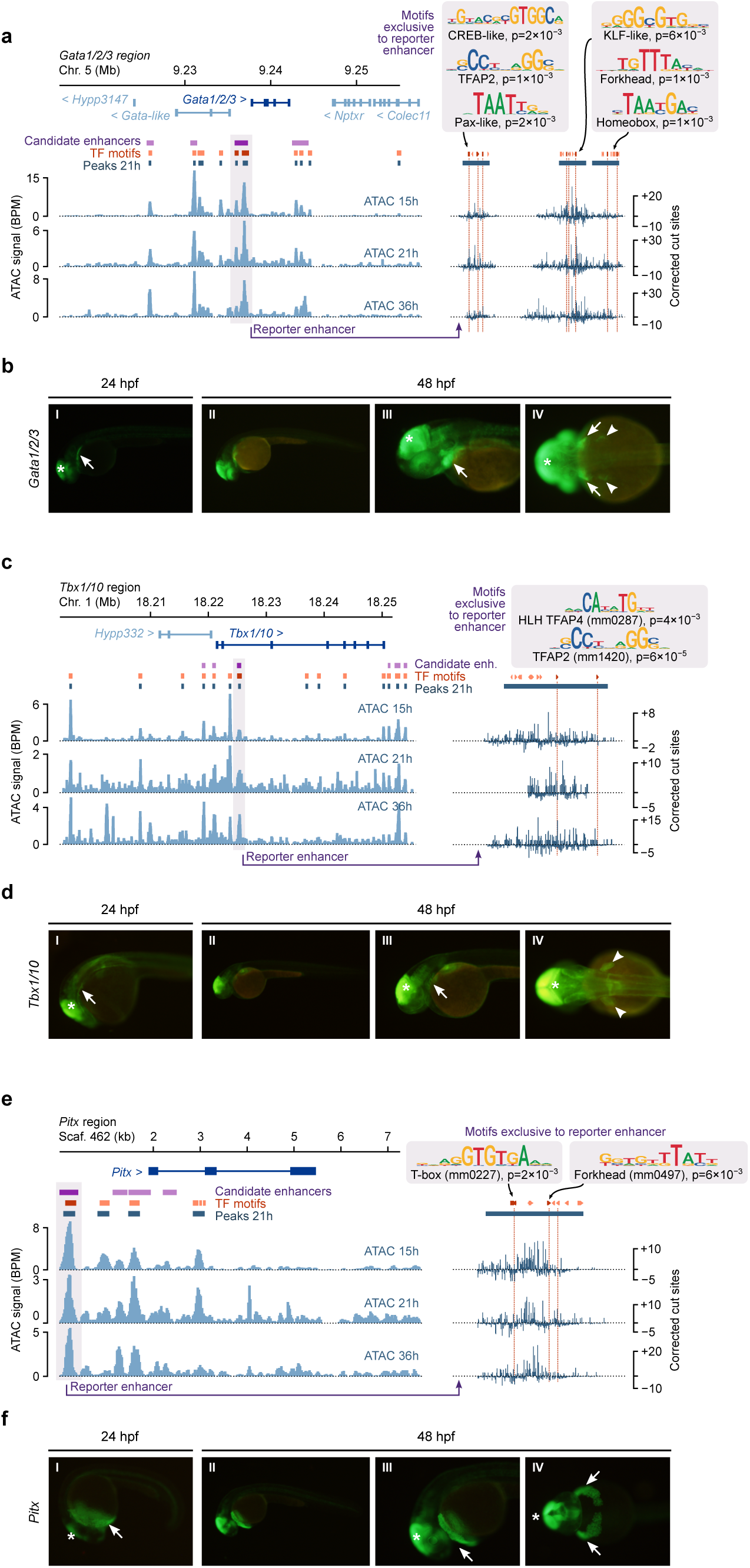
Analysis of the activity of putative regulatory regions of amphioxus genes in zebrafish. **a,** Identification of putative enhancers of *Gata1/2/3* in amphioxus, based on the examination of bulk ATAC-seq experiments (at 15 hpf, 21 hpf, and 36 hpf, measured in bins per million mapped reads, or BPM). ATAC-seq peaks at 21 hpf are showed in dark grey. Candidate enhancer regions are shown in purple. Mapped TF motifs are shown in red. The right panel to the right shows a zoom-in of the enhancer region cloned in the reporter construct in panel b (grey-shaded region), highlighting some of its unique TF motifs (top, *p*-values reflect significance of enrichment of the motif in that genomic window; see *Methods* and **Supplementary Table 2** for the complete list) and the TF binding signatures for each ATAC-seq library (expressed as *TOBIAS*-corrected ATAC cut sites, where negative values indicate regions that are putatively bound by a protein). **b**, GFP signal in F1 transgenic zebrafish embryos for the *Gata1/2/3* construct. Subpanels I to III show the lateral view (anterior to the left) of 24 hpf or 48 hpf embryos showing green fluorescence in the pharyngeal mesoderm (arrows). Subpanel IV shows a dorsal view of the same 48 hpf individual from panel III with green fluorescence in the fin buds (arrowheads). The fluorescence observed in the midbrain corresponds to the positive control included into the reporter constructs and is indicated by a white asterisk. **c,** Same as panel a, indicating putative enhancers of *Tbx1/10* in amphioxus (left) and the unique motifs and TF binding signatures of the reporter enhancer (right). **d,** Same as panel b, showing green fluorescence in the pharyngeal mesoderm from lateral viewpoints (arrows, subpanels I to III) and fin buds from a dorsal viewpoint (arrowheads, subpanel IV), at different developmental stages (24 and 48 hpf). **e,** Same as panels a and c, indicating putative enhancers of *Pitx* (left) and the unique motifs and TF binding signatures of the reporter enhancer (right). **f,** Same as panels b and d, showing green fluorescence in the hatching gland cells from lateral (arrows, subpanels I to III) and ventral (IV) viewpoints, at different developmental stages (24 and 48 hpf).

In vertebrates, both the anterior axial (prechordal plate) and pharyngeal/cranial mesoderm structures develop into different muscle populations: the extraocular muscles, and several facial/branchial muscles, respectively^10^. Interestingly, myogenesis in these cells, although it is mediated by the activity of members of the MRF family, is controlled by the upstream factors *Pitx2* (extraocular muscles) and *Tbx1* (pharyngeal muscles) and not by *Pax3/7* and *Six1/2* factors as it is the case for muscles deriving from the somites^10,88,93^. In amphioxus, we previously showed that all the somites form under the control of *Pax3/7*, *Six1/2* and/or *Zic*^11^. Moreover, *Pitx*, the ohnologue of vertebrate *Pitx1*, *Pitx2* and *Pitx3*, has been shown by *in situ* hybridization to be expressed on the left side of the embryo and in few neurons and is controlling left/right asymmetry^26,72,94^, while we observed in our data its expression only in two metacells (3 and 11) in the somite subclustering atlas (**Supplementary Fig. 5**). On the other hand, *Tbx1/10* has been shown to be expressed long after MRFs in the amphioxus somites^11,71^ and we showed in our data a reduced expression in metacell 8 in the somite subclustering atlas (**Supplementary Fig. 5**), metacell we assigned to the muscular part of the trunk somites, while its expression was not detected in the other metacells expressing MRFs. If our scenario of head mesoderm evolution is correct, it implies that *Pitx2* and *Tbx1* were co-opted for the control of myogenesis in the vertebrate head. In order to test this co-option, we searched for putative regulatory regions for both genes using ATAC-seq data and tested them in zebrafish reporter assays, as described above for *Gata1/2/3*. We cloned eight ATAC-seq peak regions around *Tbx1/10* (**Fig. 5c**), and six around *Pitx* (**Fig. 5e**) and we tested their activity by generating F0 transgenic zebrafish. In the case of Tbx1/10, only one region was able to drive the reporter gene expression (*eGFP*) in a restricted manner and we generated the corresponding F1 transgenic lines. The genomic region tested controlled the expression of the reporter gene in the zebrafish head pharyngeal mesoderm at 24 hpf and in both the head mesoderm and the finbuds at 48 hpf (**Fig. 5d**) and it contains motifs for HLH class TFs (**Fig. 5a**, **Supplementary Table 2**). Among this family of TF, several zebrafish *Twist* paralogues are expressed in both head mesoderm and pectoral fin bud^95^. This suggests that *Tbx1/10* in the chordate ancestor probably contained regulatory information that allowed its later recruitment in the vertebrate head mesoderm for a new function as a myogenesis controlling factor. In the case of *Pitx*, also only one region drove a restricted reporter expression in zebrafish at F0 and was used for generating F1 lines. In this case, expression was observed in the hatching gland at both 24 hpf and 48 hpf (**Fig. 5f**). The zebrafish hatching gland derives from the anterior prechordal plate and it expresses *Pitx2* during embryogenesis^91,96–98^. Interestingly, the putative enhancer region used in the tested construction contains a T-box class motif, potentially recognized by *Tbx16* from zebrafish, which is expressed in the prechordal plate^99,100^, and a Forkhead-class motif, potentially recognized by *Foxh1*, which is a downstream effector of the Nodal signalling pathway^101^, the nodal ligand gene *ndr2* being expressed in the zebrafish prechordal plate^102^ (**Fig. 5e, Supplementary Table 2**). Hence, our result suggests that the *Pitx* gene in the chordate ancestor already had the potentiality to be recruited in this mesoderm region during vertebrate evolution.

To conclude, our cell atlas and transgenesis approaches support a scenario for the emergence of the vertebrate lateral plate mesoderm and cranial/pharyngeal mesoderm through the segregation of pre-existing cell populations (homologous to amphioxus ventral part of the somites, first pair and posterior, respectively), which, by becoming partly independent from the somites, could evolve new structures in the trunk and in the head (**Fig. 6**). We also bring new arguments for the existence of a prechordal plate-like territory in amphioxus and give insights into how the appearance of vertebrate head muscles developing from the prechordal plate and cranial/pharyngeal mesoderm might have been achieved by the co-option of *Pitx2* and *Tbx1* for the control of myogenesis.

**Figure 6.**
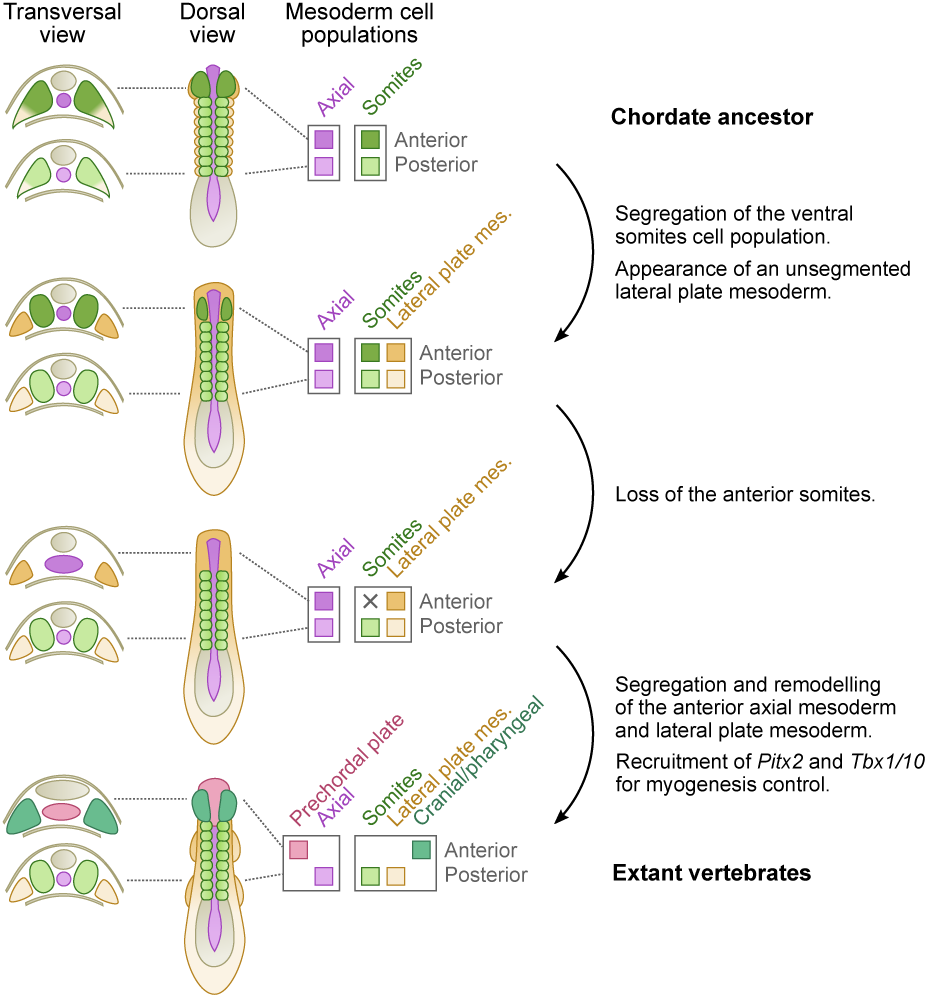
Evolutionary scenario for mesoderm evolution in chordates. Schemes of putative embryos in dorsal views with anterior to the top are shown, with transverse sections at the level of the anterior and trunk regions on the left. Diagrams on the right represent mesoderm cell populations that were inferred at each step. We propose that the chordate ancestor possessed a mesoderm organized in an axial domain with two cell populations: a prechordal-plate like region in the anterior part (dark purple), and a notochord (light purple) more posteriorly; and a paraxial domain completely segmented into somites (green), containing in the ventral part a cell population homologous to the ventral part of amphioxus somites (orange), and showing heterogeneity between the anterior (dark orange/green) and trunk (light orange/green) regions. During the first step of evolution, we propose that the ventral somite cell populations became independent from the paraxial mesoderm to give rise to the unsegmented lateral plate mesoderm (orange). In a second step, the anterior paraxial mesoderm would have been lost (dark green), and we previously proposed that this could be due to a change in the function of the FGF signalling pathway^11,12^. This loss would have led to a relaxation of the developmental constraints imposed by the segmented paraxial mesoderm in the anterior region, enabling remodelling of the tissues of the anterior axial mesoderm (dark purple) and anterior lateral mesoderm (dark orange), which could have evolved into the prechordal plate (pink) and pharyngeal/cranial mesoderm (blue/green). The ability of these new embryonic structures, derived from non-myogenic cell populations, to form muscles, would have been associated with the co-option of *Pitx2* and *Tbx1/10* as master genes of the myogenesis program.

## Material and methods

### Cell suspension preparation

Adult amphioxus (*Branchiostoma lanceolatum*) were collected at the Racou beach near Argelès-sur-Mer, France. Gametes were obtained by heat stimulation as previously described in (Fuentes, Benito et al. 2007). Embryos (∼100) at 21 hours post-fertilization (hpf, at 19°C) were washed 2 times in Ca2+/Mg2+ -free and EDTA-free artificial seawater (CMFSW : 9 mM KCl, 449 mM NaCl, 33 mM Na2SO4, 2,15 mM NaHCO3, 10 mM Tris-HCl). CMFSW was replaced by CMFSW with Liberase TM at 250µg/mL. Cells were then dissociated by a serie of pipetting and vortexing during 25 minutes at room temperature. The reaction was stopped by the addition of 1/10th volume of 500 mM EDTA. The cell suspension was centrifuged at max speed for 1 min. The pellet was resuspended in CMFSW containing Calcein violet and Propidium iodide (1 µg/mL).

### MARS-seq

Live single cells were selected using a FACSAria II cell sorter. To this end, we sorted only Calcein positive/PI negative cells, and doublet/multiplet exclusion was performed using FSC-W versus FSC-H. Cells were distributed into 384-wells capture plates containing 2 µl of lysis solution: 0.2% Triton and RNase inhibitors plus barcoded poly(T) reverse-transcription (RT) primers for single cell RNA-seq. Single cell libraries were prepared using MARS-seq^18^. First, using a Bravo automated liquid handling platform (Agilent), mRNA was converted into cDNA with an oligo containing both the unique molecule identifiers (UMIs) and cell barcodes. 0.15% PEG8000 was added to the RT reaction to increase efficiency of cDNA capture. Unused oligonucleotides were removed by Exonuclease I treatment. cDNAs were pooled (each pool representing the original 384-wells of a MARS-seq plate) and linearly amplified using T7 in vitro transcription (IVT) and the resulting RNA was fragmented and ligated to an oligo containing the pool barcode and Illumina sequences, using T4 ssDNA:RNA ligase. Finally, RNA was reverse transcribed into DNA and PCR amplified. The size distribution and concentration of the resulting libraries were calculated using a Tapestation (Agilent) and Qubit (Invitrogen). scRNA-seq libraries were pooled at equimolar concentration and sequenced to saturation (median 6 reads/UMI) on an Illumina NextSeq 500 sequencer and using high-output 75 cycles v2.5 kits (Illumina), obtaining 483M reads in total. To quantify single-cell gene expression, MARS-seq reads were first mapped onto *Branchiostoma lanceolatum* genome (GCA_927797965.1, annotation version 3) using STAR v2.7.3^103^ (with parameters: –*outFilterMultimapNmax* 20 –*outFilterMismatchNmax* 8) and associated with exonic intervals. Mapped reads were further processed and filtered as previously described^18^. Briefly, UMI filtering includes two components, one eliminating spurious UMIs resulting from synthesis and sequencing errors, and the other eliminating artefacts involving unlikely IVT product distributions that are likely a consequence of second strand synthesis or IVT errors. The minimum FDR q-value required for filtering in this study was 0.02.

### Single cell transcriptome clustering

We used Metacell 0.37^19^ to select gene features and construct high-granularity cell clusters (metacells), which were further annotated into cell types (see below). First, we selected informative genes using the *mcell_gset_filter_multi* function in the *metacell* R library, including genes fulfilling these criteria: a total gene UMI count > 30 and >2 UMI in at least three cells, a size correlation threshold of −0.1, and a normalized niche score threshold of 0.01. This resulted in the selection of 844 genes to be used for downstream clustering. Second, we used these genes to build a *K-*nearest neighbours cell graph with *K* = 100 (*mcell_add_cgraph_from_mat_bknn* function), which was the basis to define metacells with an additional *K-*nearest neighbour procedure (*mcell_coclust_from_graph_resamp* and *mcell_mc_from_coclust_balanced* functions) using *K =* 30, minimum metacell size of 15 cells, and 1,000 iterations of bootstrap resampling (at 75% of the cells); and a threshold α = 2 to remove edges with low co-clustering weights. Third, we removed one metacell which exhibited low transcriptomic information (> 50 cells with a median UMI/cell < 500). This resulted in 176 metacell clusters, which were annotated to known cell types (**Supplementary Table 4**) based on the expression level of known markers (**Extanded Data Fig. 1-6**).

We recorded gene expression in cell clusters (metacells or cell types) by computing a regularized geometric mean within each cluster and dividing this value by the median across clusters. This normalized gene expression can be interpreted as an expression fold change (FC) for a given metacell or cell type.

Two-dimensional projection of the metacells were created using a force-directed layout based on the metacell co-clustering graph (*mcell_mc2d_force_knn* function).

Gene expression profiles across cell clusters were visualized with heatmaps, using the *ComplexHeatmap* 2.10.0 R library^104^. Cell cluster ordering was fixed according to annotated cell types; and gene order was determined using the highest FC value per cluster. Genes were selected based on minimum differential expression per metacell/cell type, with a maximum number of markers per clusters selected in each case (the actual thresholds used in each heatmap are specified in the corresponding figure legends).

Finally, we selected cells belonging to the endoderm, neural and somitic metacells (**Supplementary Table 4**), and reclustered them using the same *metacell*-based approach as described for the whole dataset (except that in this case we allowed for smaller metacells, with 10 cells; (**Supplementary Table 4b-d**). The two-dimensional arrangement of the resulting metacells was curated based on the expression of cell type-specific known markers of various cell subtypes (**Supplementary Fig. 4, 5 and 6**).

### ATAC-seq library preparation

For ATAC-seq library construction, 25 embryos at the 21 hpf (19°C) were transferred in a 1.5 ml tube, in four replicates. We then followed the method described in^105^. Around 50,000 cells were used for tagmentation.

### Analysis of neurula regulatory regions

We used the ATAC-seq data from the 21 hpf embryo to build a catalogue of neurula regulatory regions. For comparison, we also used previously published^15^ ATAC-seq libraries of 15 hpf and 36 hpf embryos (the closest developmental timepoints available in that study; NCBI SRA accession numbers SRR6245277 to SRR6245279), as well as H3K4me3 ChIP-seq libraries from these same timepoints (SRA accession numbers SRR6245317 to SRR6245320). The ATAC-seq libraries corresponding to the 15, 21 and 36 hpf embryos were mapped separately to the *B. lanceolatum* genome using *bwa* 0.7.17 (*mem* algorithm^106^). The resulting BAM files were (i) filtered using *alignmentSieve* (from the *deeptools* 3.5.1 package^107^) to exclude weak alignments MAPQ > 30), (ii) corrected to shift the left and right ends of reads, to account for ATAC mapping biases (+4/−5 bp in the positive and negative strands, using the --*ATACshift* flag in *alignmentSieve*), and (iii) filtered to only include nucleosome-free alignments (*--maxFragmentLength 120* with *alignmentSieve*). Duplicated reads were marked with *biobambam2* 2.0.87^108^, coordinate-sorted, and removed to produce filtered BAM files. Then, we concatenated the BAM files stage-wise. Normalized coverage for each stage was reported as bins per million mapped reads (BPM), calculated using the *bamCoverage* tool in *deeptools*. The ChIP-seq libraries for 15 and 36 hpf were processed in the same way (except for the ATAC mapping bias correction step and the filtering of nucleosome-free alignments).

For the 21 hpf ATAC-seq experiment, we used *MACS2* 2.2.7.1^109^ to identify regulatory elements with the *callpeak* utility, starting from the nucleosome-free filtered BAM file, with the following options: (i) an effective genome size equal to the ungapped amphioxus genome length, (ii) keeping duplicates from different libraries (*--keep-dup all* flag), (iii) retaining peaks with a *q-*value < 0.01, (iv) enabling multiple summit detection (*--call-summits* flag), and (v) disabling the modelling of peak extension for ChIP-seq libraries (*--nomodel* flag).

We then assigned the *MACS2*-predicted regulatory elements to their proximal genes, based on their distance to each gene’s transcription start site (TSS). Specifically, we selected well-supported *MACS2* regulatory elements (*q-*value < 1×10^−6^), standardized their lengths to 250 bp (125 bp to each side of the predicted peak summit), and assigned each peak to nearby genes based on distance to their TSS (excluding genes further away than 20 kbp, and genes located beyond a more proximal gene). Peaks overlapping the promoter region of a particular gene (defined based on TSS coordinates +/– 50/200 bp or coincidence with H3K4me3 ChIP-seq peaks for the 15 and 36 hpf datasets) were not assigned to any other gene. The peak sets were reduced to non-overlapping sets to avoid redundant regions. These genome coordinate operations were done using the *GenomicRanges* 1.46 and *IRanges* 2.28 packages in *R*^110^. We used these gene-regulatory element assignments to define lists of cell type-specific regulatory elements, based on the expression specificity of each gene (expression fold change ≥ 1.5 in a given cell type). In parallel, we also defined a set of background regulatory regions for each cell type (consistent of regulatory regions linked to non-overexpressed genes, at fold change ≤ 1). In total, we assigned 51,028 regulatory regions (ATAC peaks) to 19,069 genes (out of 27,102), with a median of 2 peaks per gene.

We used the cell type-specific sets of active regulatory elements (and their corresponding background sets) to identify motifs *de novo* using the *findMotifsGenome.pl* utility in *homer* 4.11^111^ Specifically, we set a constant peak size of 250 bp and attempted to identify motifs for each cell type, using *k*-mers of length 8, 10, 12, and 14; and tolerating up to four mismatches in the global optimization step.

In order to build a final motif collection for amphioxus, we concatenated the cell type-specific *de novo* motifs with known TF binding motifs from the CIS-BP database (as available the 3rd of March, 2023)^112^. Specifically, we used 3,547 experimentally determined motifs (with SELEX or PBMs), corresponding to vertebrate or tunicate species (*Homo sapiens*, *Mus musculus*, *Xenopus tropicalis*, *Xenopus laevis*, *Danio rerio*, *Tetraodon nigroviridis*, *Meleagris gallopavo*, *Gallus gallus*, *Anolis carolinensis*, *Takifugu rubripes*, *Ciona intestinalis*, and *Oikopleura dioica*). We reduced the redundancy of this extensive *de novo* + known motif collection based on motif-motif sequence similarity, as follows: (i) we removed motifs with *homer* enrichment *p*-values < 1×10^−9^; (ii) we retained with high contiguous information content (IC), defined as having IC ≥ 0.5 for at least four consecutive bases or IC ≥ 0.5 for two or more blocks of at least three bases; (iv) for each of the remaining motifs, we measured their pairwise sequence similarity by calculating the weighted Pearson correlation coefficient of the position probability matrices of each motif, using the *merge_similar* function in the *universalmotif* 1.12.4 (Tremblay 2022) R library with a similarity threshold = 0.95 for hierarchical clustering and a minimum overlap of 6 bp between two motifs in the motif alignment step. Finally, we selected the best motif per cluster based on its IC (highest). This resulted in a final, non-redundant collection of 1,595 motifs.

Then, we calculated the enrichment of each motif among the sets of regulatory regions specific to each cell type. To that end, we used the *calcBinnedMotifEnrR* function in the *monalisa* 1.0 R library^113^ to count motif occurrences in three sets of regulatory regions (bins) defined based on the expression levels of their associated genes: highly cell type-specific genes (FC ≥ 1.5), mildly cell type-specific genes (FC ≥ 1.1 and < 1.5), and non-cell type-specific genes (FC < 1). Motif occurrences were defined as motif alignments with scores above 80% of that motif’s maximum alignment score (defined from the corresponding position weight matrices). Motif enrichment in each bin was then calculated using the fold change of occurrence relative to randomly sampled genomic regions (matched by GC content and length, using twice as many regions for background as for the foreground), and its significance assessed using a binomial test followed by Benjamini-Hochberg *p-*value adjustment. We retained the fold change and *p*-values for th set of highly cell type-specific regulatory regions (i.e. from genes with FC ≥ 1.5) for further analysis (**Fig. 3 and Supplementary Table 1**).

Finally, we scanned the *B. lanceolatum* genome to identify discrete occurrences of each of the 1,595 motifs across the 51,028 *MACS2*-defined regulatory regions. We used the *findMotifHits* function in *monalisa.* In order to define *bona fide* motif alignments, we calculated an empirical *p-*value for each motif alignment (only best alignment per regulatory region) based on the rank of its alignment score when compared to a background distribution of randomly sampled genomic regions of similar sequence composition (only best alignment score per random background bin). Specifically, we divided the foreground regions into 10 equal-size sets based on their GC content, and matched each set with random genomic background sequences (not in the foreground) of similar GC content (same category) and equal length (set to 250 bp). These motif aligments were used to identify enhancer-specific motifs in **Fig. 5** (complete list in **Supplementary Table 2**).

### Cross-species cell type comparison

We used SAMap 1.0.2 to evaluate the similarity between *B. lanceolatum* cell types and the previously published developmental single-cell transcriptomes of *Danio rerio*^41^ (reference gene set in original study: GRCz10 v1), *Xenopus tropicalis*^40^ (reference gene set in original study: Xenbase version 9.0), and *Ciona intestinalis*^39^ (reference gene set in original study: KH2012 from the Ghost Database (http://ghost.zool.kyoto-u.ac.jp/download_kh.html).

For each query species, we used the UMI tables corresponding to the timepoints closest to the *B. lanceolatum* 21hpf developmental stage (12 in total): 14 hpf, 18 hpf and 24 hpf for *D. rerio* (GEO accession: GSE112294); S14, S16, S18, S20 and S22 for *X. tropicalis* (GSE113074); and the initial, early, middle and late tailbud stages for *C. intestinalis* (GSE131155). For *C. intestinalis*, we used the cell type annotations used in the original paper. For the two vertebrates, we used the consensus cell annotations employed by Tarashansky *et al.*^114^.

To run SAMAp, we first created a database of pairwise alignments with *blastp* 2.5.0 (comparing *B. lanceolatum* peptides to each query species separately; in the case of *Danio rerio* we used *blastx*/*tblastn* instead of *blastp* as the original gene set^41^ was only available as un-translated transcripts). Second, we used the cell-level UMI counts of each gene to calculate the SAMap mapping scores for each pair of cell types (between *B. lanceolatum* and each of the 12 query developmental datasets in other species), using all cells within each cluster for score calculation.

Finally, we identified shared marker genes between cell types of *B. lanceolatum* and the query chordate species by identifying sets of cell type-overexpressed genes with the *scanpy* 1.9.3^115^ *rank_genes_groups* function to calculate cell type-level fold change values and overexpression significance (Wilcoxon rank-sum tests followed by BH *p-*value adjustment). For each species, cell type-specific genes were then determined based on fold change and overexpression significance (at adjusted *p* < 0.05 and FC ≥ 1). For cross-speices comparisons, genes were linked based on shared orthology group membership. Orthology groups between genes of the the four species were determined using *Broccoli* 1.1^116^ (using predicted peptides as input; disabling the *k-*mer clustering step; using up to 10 hits per species for maximum-likelihood phylogenetic tree calculations; and adding two additional chordates for better coverage: *Mus musculus* and *B. floridae*).

We also performed a more detailed analysis of shared TFs between amphioxus and the other three chordates, selecting cell type-specific amphioxus TFs (*p <* 0.05 and FC ≥ 1.25; see below details on TF annotation) and evaluating whether their orthologs in chordates were also over-expressed in cell types homologous to the amphioxus endoderm (in this case, it was compared to endodermal tissues in the other chordates), endostyle (to other endodermal tissues), muscular somites (to vertebrate skeletal muscle and tunicate muscle/heart), somites (to vertebrate presomitic mesoderm or tunicate muscle/heart), notochord (to other notochordal tissues) hypothalamus and neurons (each of which was compared to vertebrate neurons, hindbrain, forebrain/midbrain, notoplate and neuroendocrine cells; and to the tunicate nervous system), and the anterior and posterior epidermis (each compared to epidermal progenitors, ionocytes, small secretory epidermal cells, goblet cells, and hatching gland).

### Gene family annotation

We ran gene phylogenies to refine the orthology assignments of TF gene families. We used translated peptide sequences from 32 metazoan (longest isoforms per gene, **Supplementary Table 3**, which were scanned using *hmmsearch* (*HMMER* 3.3.2^117^) to identify hits of TF-specific HMM profiles (from Pfam 33.0^118^) representing their corresponding DNA-binding regions. For each gene family, the collection of homologous proteins was aligned to itself using *diamond blastp* v0.9.36^119^ and clustered into low-granularity homology groups using the Markov Cluster Algorithm *MCL* v14.137^120^ (using alignment bit-scores as weights, and a gene family-specific inflation parameter; **Supplementary Table 3b**). Then, each homology group was aligned using *mafft* 7.475^121^ (E-INS-i mode, up to 10,000 refinement iterations). The alignments were trimmed with *clipkit* 1.1.3^122^ (*kpic-gappy* mode and a gap threshold = 0.7) and used to build phylogenetic trees with *IQ-TREE* v2.1^123^ (running each tree for up to 10,000 iterations until convergence threshold of 0.999 is met for 200 generations; the best-fitting evolutionray model was selected with *ModelFinder*^124^; statistical supports were obtained using the UFBoot procedure with 1,000 iterations (Hoang, Chernomor et al. 2018)). Outlier genes were removed from each tree using *treeshrink* v1.3.363 (gene-wise mode using the centroid rooting algorithm; scaling factors set to *a* = 10 and *b* = 1); and the trees were recalculated if necessary if any outgroup needed to be removed. Finally, we used *Possvm* 1.1^125^ to identify orthology groups from each gene tree (with up to 10 steps of iterative gene tree rooting), and annotated the orthogroups and the *B. lanceolatum* TFs with reference human gene names.

For genes used to assign metacells to known amphioxus embryonic territories and named in the manuscript, we either used the previously published amphioxus gene names when they exist, or a name based on fine orthology analysis. Amino acid sequences from *B. lanceolatum* were used to search Genbank for putative homologues by *blasp*. Sequences were aligned using *ClustalX*^126^. Alignments were manually corrected in *SeaView*^127^. Maximum Likelihood phylogenetic trees were reconstructed using *IQ-TREE* v2.1^123^ with default parameters (fast bootstraping and automatic best model search). Genes with no clear orthology signal were named based on the presence of known protein domains.

### In situ hybridization

DIG labeled probes were synthesized from fragments cloned into pBKS, or from PCR amplified DNA fragments purchased at IDT, using the appropriate RNA polymerase (T7, T3 or SP6) and the DIG-labeling Mix (Roche). Embryos at 21 hpf (19°C) were fixed in paraformaldehyde (PFA) 4% in MOPS buffer, dehydrated in 70% ethanol and kept at −20°C. *In situ* hybridization was undertaken as previously described in^26^. The accession numbers/sequences used for probe synthesis are given in **Supplementary Table 5**.

### Zebrafish transgenesis

The putative regulatory regions were cloned after PCR amplification on genomic DNA in the PCR8/GW/TOPO vector (Life Technologies). Using Gateway technology (Life Technologies), the inserts were then shuttled into an enhancer detection vector composed of a *gata2* minimal promoter, an enhanced GFP reporter gene, and a strong midbrain enhancer (z48) that works as an internal control for transgenesis in zebrafish^128^. Transgenic embryos were generated using the Tol2 transposase system^129^. Briefly, 1-cell stage embryos were injected with 2 nl of a mix containing 25 ng/µL of Tol2 transposase mRNA, 20ng/µL of purified vector, and 0,05% of phenol red. Injected embryos were raised until the desired stage, visualized under an Olympus SZX16 fluorescence stereoscope and photographed with an Olympus DP71 camera.

### Statement that all experiments were performed in accordance with relevant guidelines and regulations

All the experiments were performed following the Directive 2010/63/EU of the European parliament and of the council of 22 September 2010 on the protection of animals used for scientific purposes. Ripe adults from the Mediterranean invertebrate amphioxus species (*B. lanceolatum*) were collected at the Racou beach near Argelès-sur-Mer, France, (latitude 42° 32′ 53′ ′ N and longitude 3° 3′ 27′ ′ E) with specific permission from the Prefect of Region Provence Alpes Côte d’Azur. Zebrafish embryos were obtained from AB and Tübingen strains, and manipulated following protocols approved by the Ethics Committee of the Andalusia Government and the national and European regulation established.

## Data availability

Accession numbers of sequences used for in situ hybridization probe synthesis are given in Supplementary Tables 5. The accession numbers for the sequences are available in Genbank.

## Supplementary Files

**Supplementary Figure 1. scRNA-seq and ATAC-seq summary statistics (related to Fig. 1 and 3). a,** Number of cells per metacell cluster. distribution of UMIs/cell in each metacell, and total number of UMIs per metacell. **b,** Fraction of reads in each ATAC-seq sample (and the pooled dataset) that are duplicated, nucleosome-free (NFR), or mapping in peaks. **c,** Inter-sample similarity for the ATAC-seq replicates, measured using the Pearson correlation coefficient of binned raw counts in the nucleosome-free fraction (bin size = 10 kbp). **d,** Insert size distribution of the pool of ATAC-seq replicates. The dotted line indicates the threshold to define the nucleosome-free fraction (120 bp). **e,** Enrichment of ATAC-seq signal around transcription start sites (TSS), calculated using binned normalised coverage (bin size = 50 bp). **f,** Fraction of ATAC-seq peaks overlapping various features in the genome. **g,** Cumulative distribution of the normalised ATAC-seq signal at the TSS of genes, sorted in five equally-sized bins according to their expression levels (low to high, measured in UMI counts). Highly expressed genes in our scRNA-seq data exhibit stronger bulk ATAC-seq signals. **h,** Distribution of number of ATAC-seq peaks detected per gene, in global (left) and for specific subsets of gene families (TFs, signalling-related genes, and neural-related genes; right).

**Supplementary Figure 2. *In situ* hybridization of genes showing an enriched expression in some metacells and for which expression was not previously described**. *In situ* hybridization experiments were undertaken on N3 stage embryos. Dorsal views with anterior to the left (top) and side views with anterior to the left and dorsal to the top are shown for each gene. Schemes of embryo showing in blue the region in which each series of genes is expressed is presented on the left. Below each *in situ* hybridization picture, transcriptomic expression of the marker is shown as density maps representing UMI counts (per 10,000 UMIs) in each cell, using the same two-dimensional metacell arrangement as in Fig. 1.

**Supplementary Figure 3. Transcription factor expression and motif activity (related to Fig. 1 and Fig. 3). a,** Normalized fold change expression of top variable TFs (rows) per metacell (columns, grouped by cell type). For each metacell, we selected TFs with a minimum fold change ≥ 2 and a total of 10 UMIs across all cells. Gene names in bold indicate that the gene is mentioned in the manuscript. **b,** Enrichment fold change of top variable TF binding motifs (rows) per cell type (columns). For each cell type, we selected up to 60 motifs with a minimum fold change ≥ 1.2 and enrichment BH-adjusted *p-*value < 0.05. Motifs are color-coded based on their sequence similarity to motifs of known TF structural classes (see *Methods*): light gray indicates *de novo* motifs without similar motifs in known databases, whereas dark gray and other colors indicate motifs that can be mapped to one or more previously described TF binding motifs. Motifs in bold are mentioned in the manuscript.

**Supplementary Figure 4. Gene expression distribution on 2D projected cells for neural gene markers (related to Fig. 3). a,** Schematics of inferred neural metacell locations over an amphioxus neurula-stage embryo, dorsal view. **b,** Gene expression is shown as density maps representing UMI counts (per 10,000 UMIs) in each cell, with cells positioned in the vicinity of their corresponding metacells. The metacells have been arranged based on their inferred position in the neurula embryo (Fig. 3a). Markers were selected from the litterature and from ISH analysis of newly discovered genes overexpressed in specific metacells in our dataset.

**Supplementary Figure 5. Gene expression distribution on 2D projected cells for endoderm gene markers (related to Fig. 3). a,** Schematics of inferred endoderm metacell locations over an amphioxus neurula-stage embryo, lateral view. **b,** Gene expression is shown as density maps representing UMI counts (per 10,000 UMIs) in each cell, with cells positioned in the vicinity of their corresponding metacells. The metacells have been arranged based on their inferred position in the neurula embryo (Fig. 3c). Markers were selected from the litterature and from ISH analysis of newly discovered genes overexpressed in specific metacells in our dataset.

**Supplementary Figure 6. Gene expression distribution on 2D projected cells for somite gene markers (related to Fig. 3). a,** Schematics of inferred somite metacell locations over an amphioxus neurula-stage embryo, side view. **b,** Gene expression is shown as density maps representing UMI counts (per 10,000 UMIs) in each cell, with cells positioned in the vicinity of their corresponding metacells. The metacells have been arranged based on their inferred position in the neurula embryo (Fig. 3e). Markers were selected from the litterature and from ISH analysis of newly discovered genes overexpressed in specific metacells in our dataset.

**Supplementary Table 1. Shared orthologous markers between chordate developmental cell types and stages (related to Fig. 2).** This table includes genes overexpressed in various cell types of the *B. lanceolatum* neurula transcriptome (reference species 1) and their overexpressed orthologs in other species (species 2 column: *C. intestinalis* or Cint, *X. tropicalis* or Xentro, or *D. rerio* or Drer). For each gene pair, we indicate in which cell type it is overexpressed in each species (and, for the query species 2, which developmental stage); their expression fold change and BH-adjusted enrichment *p-*value (from Wilcoxon rank sum tests) in both species; the gene name in amphioxus; and whether the gene is a TF or not.

**Supplementary Table 2. TF binding motifs in *Tbx1/10*, *Gata1/2/3* and *Pitx* candidate enhancers (related to Fig. 5).** List of motifs aligned to each ATAC-seq peak in the vicinity of three regions of interest around the TFs *Tbx1/10*, *Gata1/2/3* and *Pitx*. For each aligned motif, we list the regulatory region where it was found, whether the region was found to drive specific expression in zebrafish embryos (“is expression driver” column; Fig. 5), whether the region was tested in zebrafish (“is cloned” column), the motif ID and its annotation based on similarity to known CIS-BP motifs; whether the motif is exclusive to the regulatory region found to drive expression (“is motif exclusive to successful driver?” column), its alignment coordinates along the genome, its alignment score and empirical *p*-value, and the aligned sequence.

**Supplementary Table 3. Gene family annotation information. a,** Species used for the gene phylogenetic analyses of TFs, including the data sources and their taxonomy. **b,** List of TF families analyzed, including the representative Pfam domains, the *hmmsearch* threshold strategy, and the inflation parameter employed in MCL clustering. **c,** Phylogeny-based classification of amphioxus TFs, with orthogroup names taken from the human orthologs of each gene (using *Possvm*).

**Supplementary Table 4. Cell type annotation table. a,** Cell type annotations of all metacell clusters in the neurula transcriptome. For each metacell, we indicate its cell type, developmental layer, and whether it has been included in further reclustering analyses. **b-d,** Annotations of metacells for the neural, endodermal and somitic reclustering analyses.

**Supplementary Table 5**. Sequences used for probe synthesis.

## Acknowledgements

This work benefited from access to the Observatoire Océanologique de Banyuls-sur-Mer, an EMBRC-France and EMBRC-ERIC site. Embryo imaging experiments were undertaken using the material of the BIOPIC platform. The laboratory of H.E. and S.B. was supported by the CNRS, and by the “Agence Nationale de la Recheche” under the grants ANR-19-CE13-0011 and ANR-21-CE13-0034. Research in A.S-P. group was supported by the European Research Council (ERC-StG 851647) and the Spanish Ministry of Science and Innovation (PID2021-124757NB-I00). X.G-B. is supported by the European Union’s H2020 research and innovation program under Marie Sklodowska-Curie grant agreement 101031767. A.E. is supported by FPI PhD fellowships from the Spanish Ministry of Science and Innovation. J.J.T. was supported by the Spanish Ministerio de Economía y Competitividad (grant PID2019-103921GB-I00). M.I. laboratory research has been funded by the European Research Council (ERC) under the European Union’s Horizon 2020 research and innovation program (ERCCoG-LS2-101002275 to MI), by the Spanish Ministry of Economy and Competitiveness (PID2020-115040GB-I00 to MI) and by the ‘Centro de Excelencia Severo Ochoa 2013-2017’(SEV-2012-0208).

## Author contributions

Conceptualization of this study was done by J.L. G-Z., M.I., S.B., A.S.P.and H.E.; the study was carried out by X.G.B., L.S., L.M., A.S, A.N., O.F., M.I., S. B., A.S.P. and H.E.; writing of the original draft was done by X.G.B., S.B., A.S.P. and H.E.; funding was acquired by A.S.B., J.T., M.I., S.B., A.S.P. and H.E.; this study was supervised by J.L. G-Z, J.T., M.I., S.B., A.S.P. and H.E.

