## Supplementary figures and images for "An amphioxus neurula stage cell atlas supports a complex scenario for the emergence of vertebrate head mesoderm"

### Supplementary Figure 1

##### Extended Data Figure 1

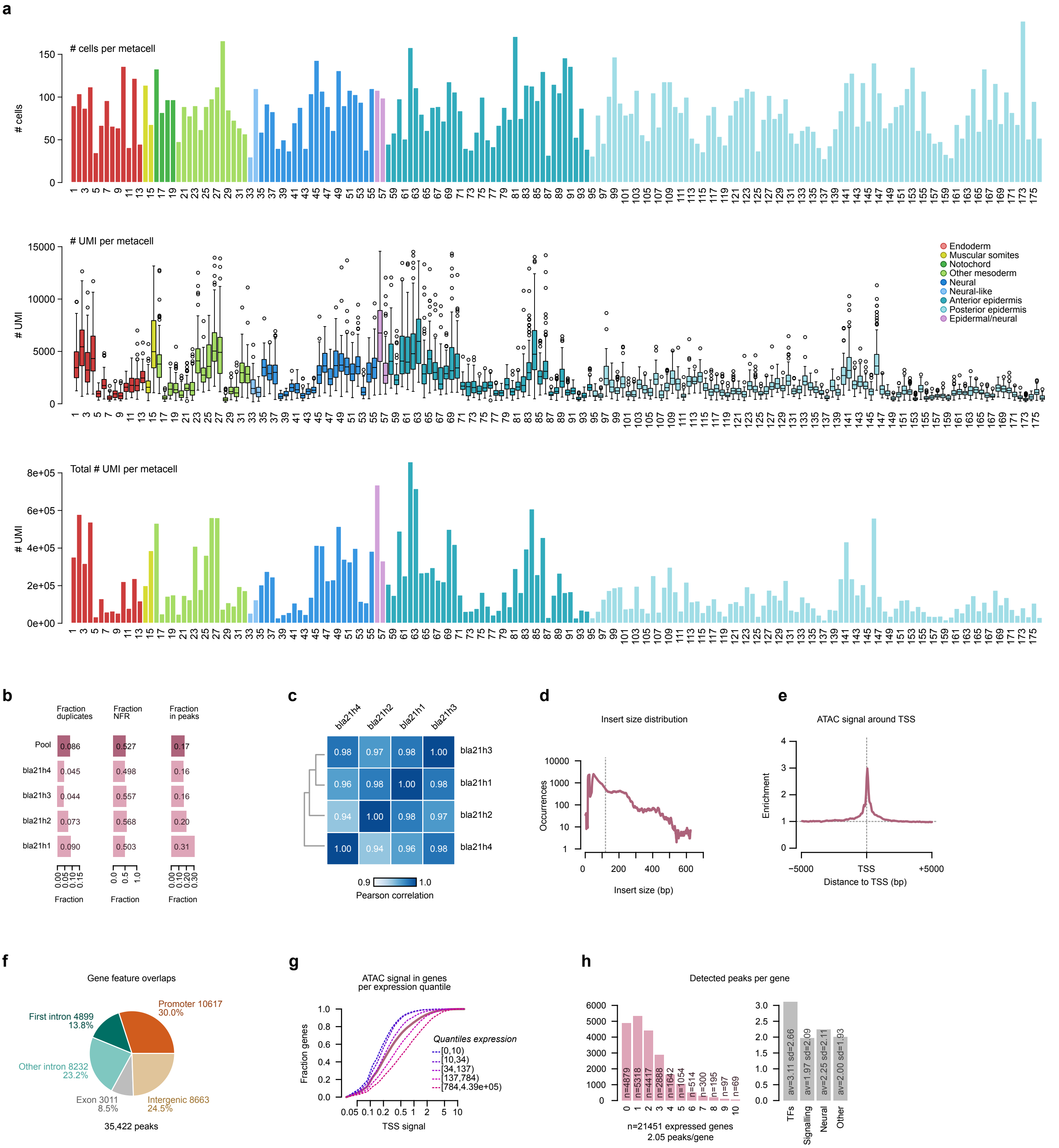

### Supplementary Figure 2

**Extended Data Figure 2**

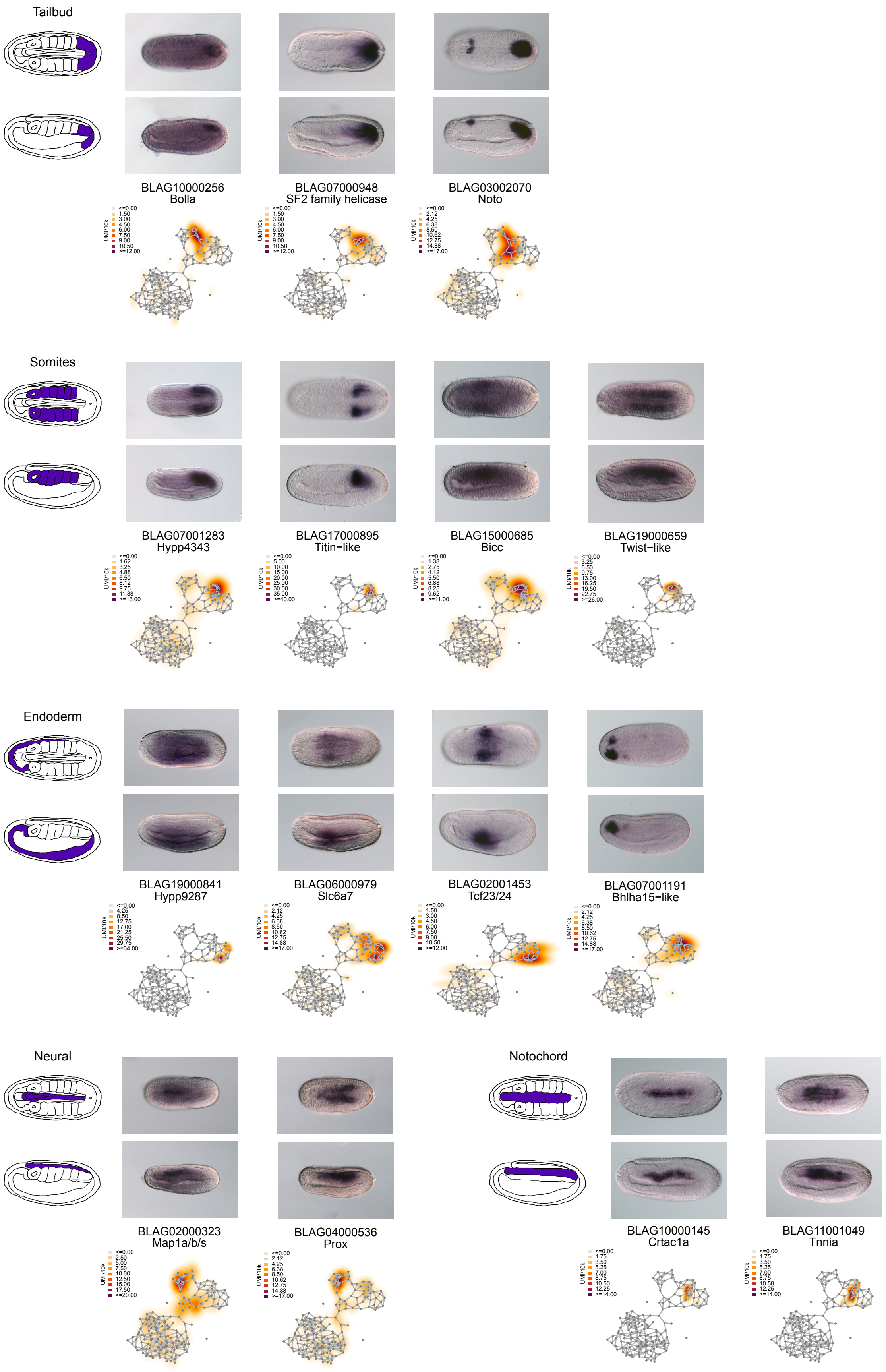

### Supplementary Figure 3

#### Extended Data Figure 3

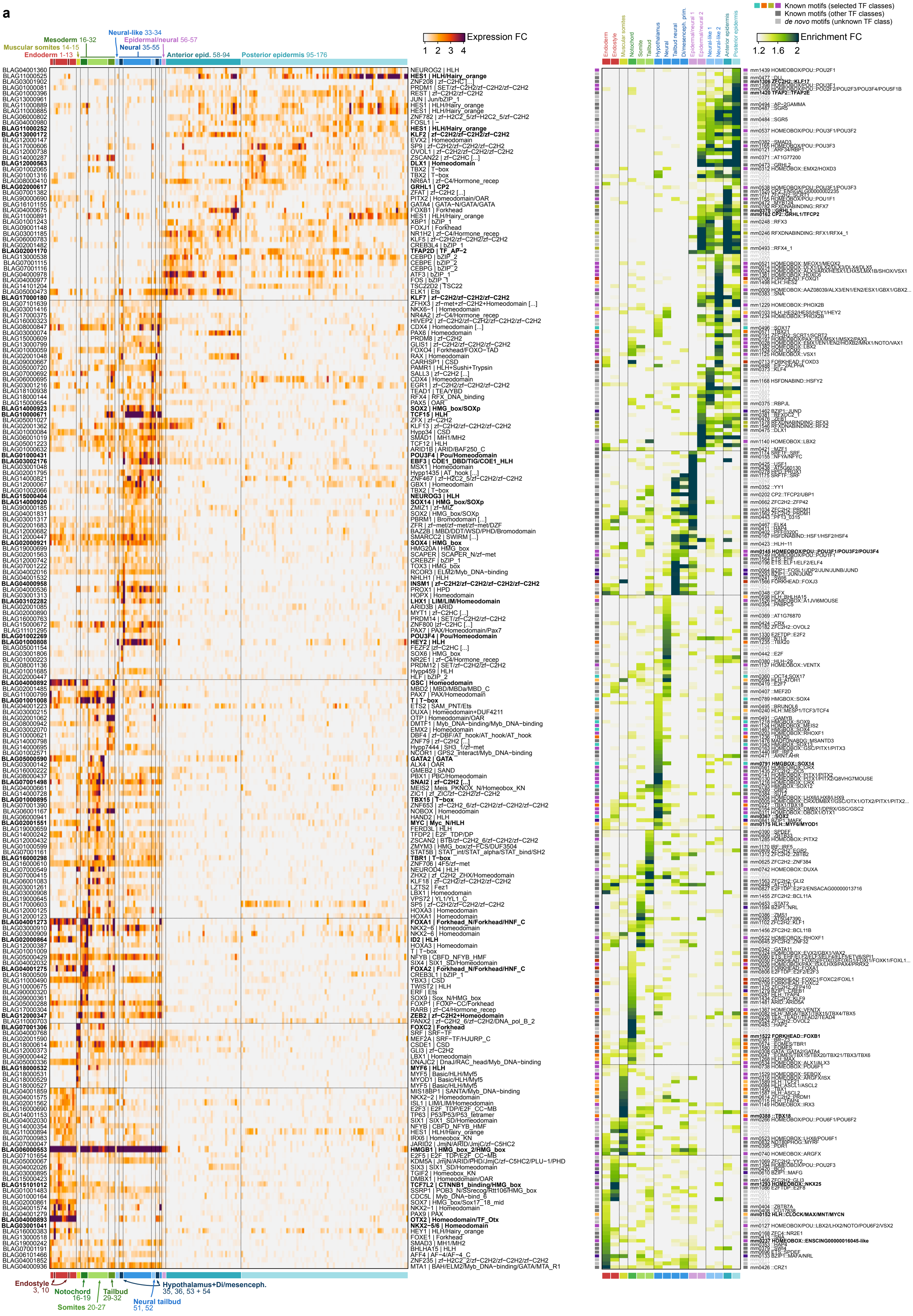

### Supplementary Figure 4

##### Extended Data Figure 4

**a**

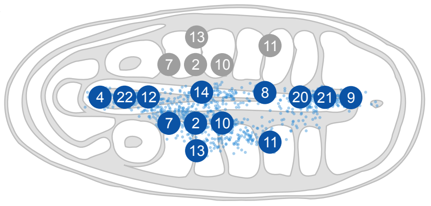


**b**

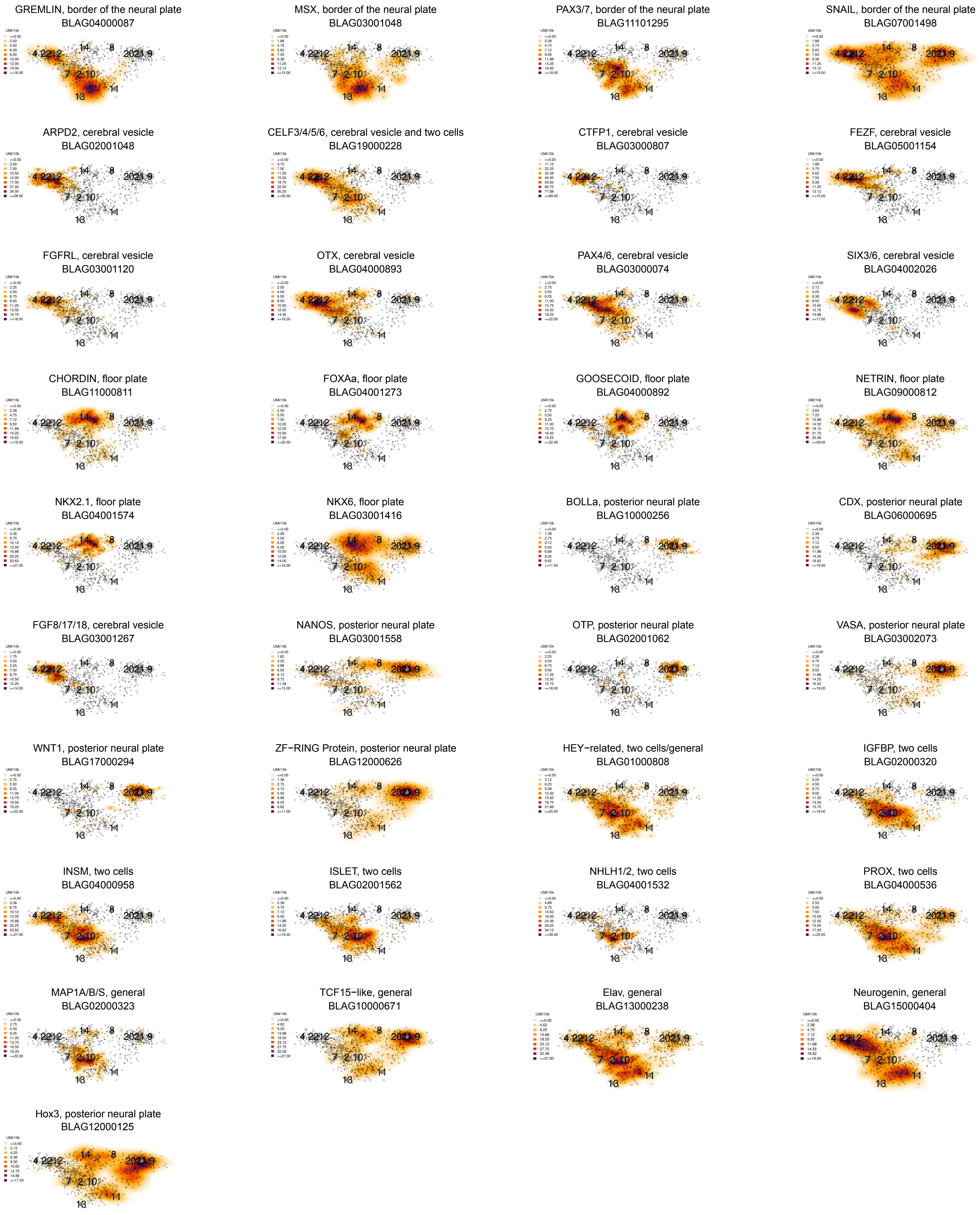

### Supplementary Figure 5

##### Extended Data Figure 5

**a**

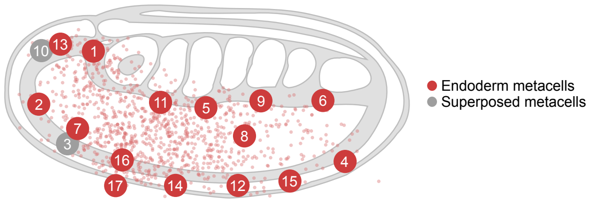


**b**

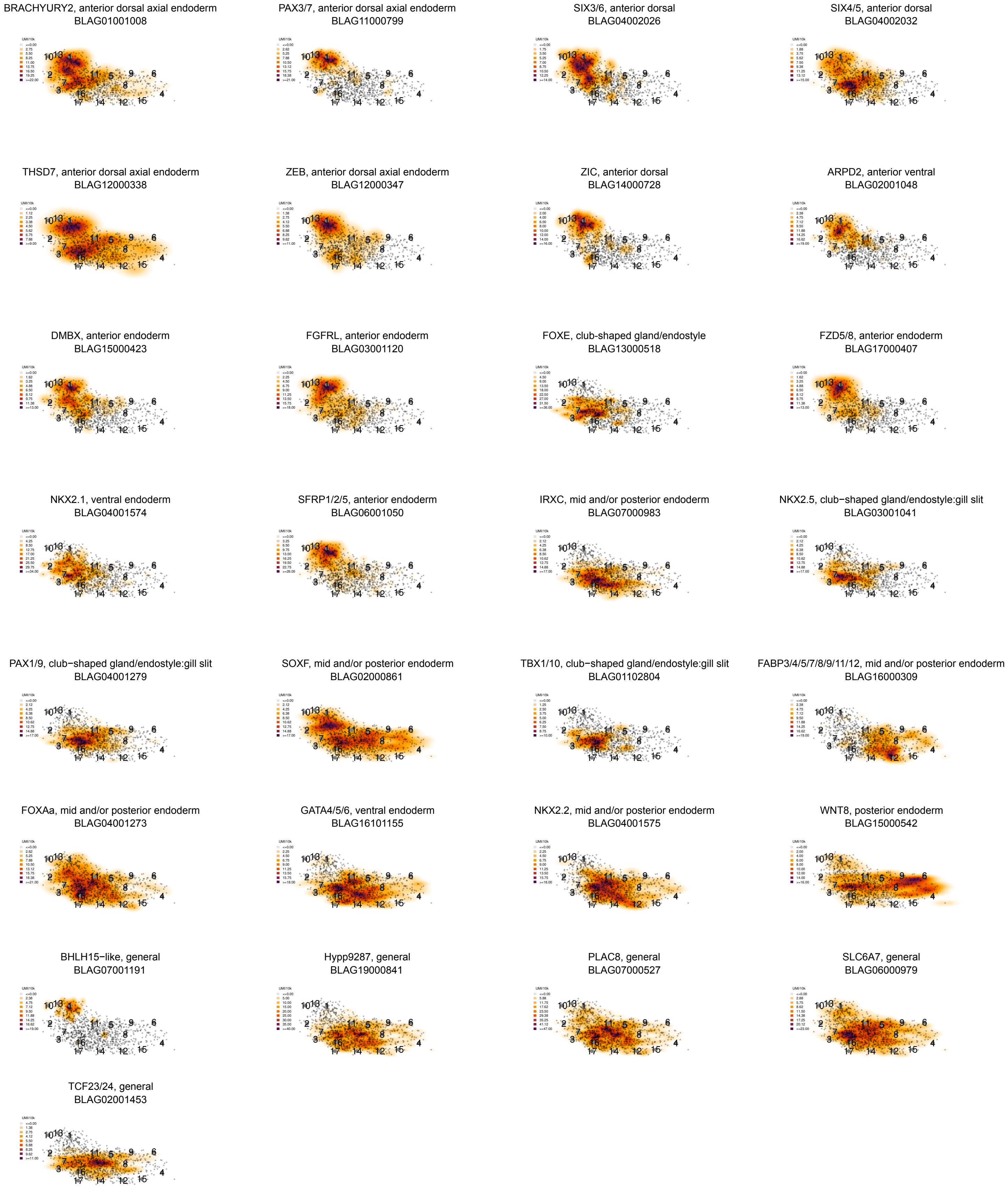

### Supplementary Figure 6

##### Extended Data Figure 6

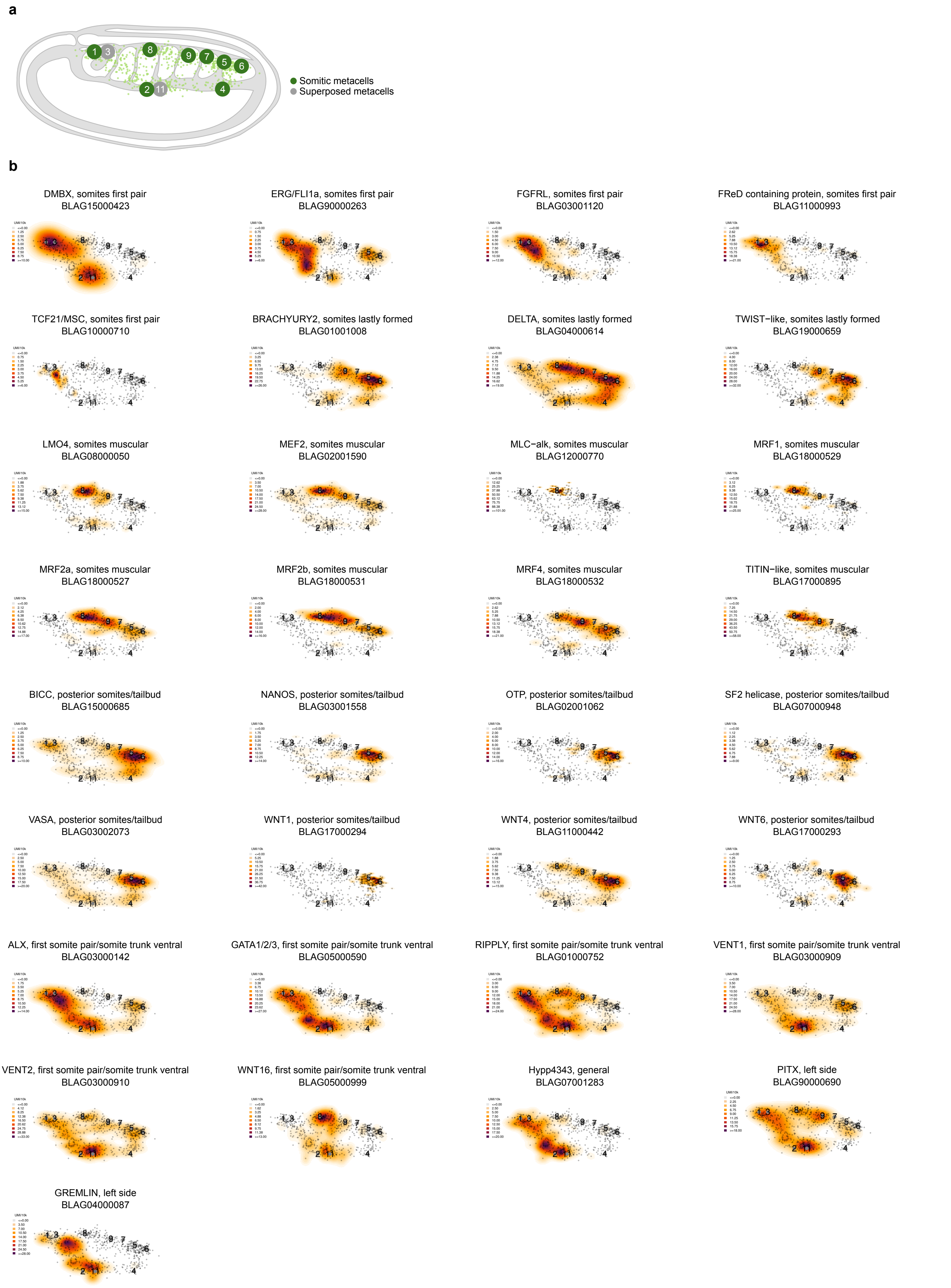
